# Predicting the alternative conformation of a known protein structure based on the distance map of AlphaFold2

**DOI:** 10.1101/2024.06.09.598121

**Authors:** Jiaxuan Li, Zefeng Zhu, Chen Song

## Abstract

With AlphaFold2 (AF2) becoming the top structural prediction tool, multiple studies have found that AF2 often favors one conformation state over others in high-precision structure predictions. Meanwhile, it has also been demonstrated that the prediction of multi-state structures from a given protein sequence is possible by subsampling multiple sequence alignment (MSA). In this work, we reveal that AF2 predictions contain information on multi-state structures even with the deepest MSA: protein distance maps extracted from AF2 often exhibit multi-peak signals in the distance probability distributions for residue pairs. By fitting and separating these multi-peak distributions of residue pairs, one can extract distinct distance information of two states, which can be incorporated into Rosetta as restraint energy functions to model large and complex conformational changes. Twenty protein systems with different types of conformational changes were selected for validation in modeling their alternative conformations. With our protocol, we successfully predicted the alternative conformations of 19 systems and achieved a template-based modeling score (TM-score) above 0.90 for the best-sampled models in nine cases. This work further expands the usage of AlphaFold2 in studying multi-state proteins.

## 1 Introduction

Proteins present versatile functional conformation states to execute the biological activities essential for living systems. ^1,2^ Modeling protein structures *in silico* has been a pivotal biological challenge studied for decades. Thanks to advancements in deep learning, accurately predicting protein structures from sequences has become commonplace. Presently, coevolution-based techniques represent the cutting edge, with AlphaFold2^3^ (AF2) standing among the foremost prediction models. The state-of-the-art methods, such as AF2, RoseTTAFold, and trRosettaX, etc. have significantly improved prediction accuracy of protein structures, often reaching an accuracy comparable to experimental results. ^4–6^ These deep-learning-based methods also facilitated other related studies, such as utilizing predicted structures as AI-augmented inputs for molecular dynamics (MD) simulations. ^7,8^

Furthermore, the advent of AF2 has sparked renewed inspiration in predicting multi-state conformations, which is crucial for comprehending dynamic behaviors and exploring the conformation space from the structural biology perspective. ^9–12^ Nonetheless, determining multiple functional states of proteins, such as the apo and holo state of enzymes and the outward/inward-facing states of transporter proteins, remains challenging both experimentally and computationally. Current prediction methods, AF2 included, primarily concentrate on predicting one particular state. ^13^ Predicting alternative conformations and assessing the impact of sequence mutations still pose difficulties with standard AF2 usage. ^14,15^ In CASP15, prediction targets encompassed ensembles of protein conformations, ^16^ and there is growing interest in developing computational approaches to predict alternative conformations in the field.

There have been many attempts targeting multi-state conformation predictions through AF2 with certain modifications. It is realized that one can further increase AF2’s sampling diversity via enabling the dropout layers and switching random seeds. ^17^ However, this approach only contributes subtle changes to the default predictions. Manipulation of multiple sequence alignment (MSA) offers great potential for predicting multiple conformations. The prevalent strategy involves MSA-subsampling, as reducing MSA depth can introduce greater uncertainty into AF2’s predictions. ^18,19^ It has effectively sampled alternative conformational states of transporters by randomly subsampling MSAs down to just 16 sequences. ^18^ However, this line of work may compromise prediction quality or heavily rely on known structural templates. ^18^ Notably, Wayment-Steele et al. successfully extracted the evolutionary signals from different states of metamorphic proteins by clustering the MSAs. ^20^ Replacing the homology search algorithm and increasing the quality of homologous sequences also contribute to the prediction of relative state populations. ^21^ The sensitivity between AF2 samples and MSAs can be further explored through sequence mutation analysis ^22^ or Boltzmann weights. ^7,23^ More interestingly, several recent deep-learning-based methods used generative models to learn the distribution of multiple conformations, ^24,25^ being a promising direction. Apart from these, another line of work has successfully incorporated explicit ligand conditions to predict the holo-states of proteins. ^26–28^

In a recent study, we demonstrated that deep learning-based models can *de novo* capture multistate information of a protein structure, and proposed a protocol to model the alternative conformation from a known structure based on contact map predictions. ^29^ In this paper, we further explore multi-conformation prediction methods by using AF2 and predominantly focus on predicting the alternative conformations of two-state proteins. We investigate the possibility of accessing these alternative conformations directly from AF2 predictions, without resorting to MSA depth reduction. Starting from a known structure, we incorporated restraints derived from distance maps of AF2 and successfully achieved satisfactory predictions of the alternative state for 20 selected two-state proteins.

## 2 Results

### 2.1 AlphaFold2 predictions with full MSA contain alternative conformational information

Our study examined 20 proteins known to exist in two distinct structural states. These proteins range in size from 90 to 600 amino acids and encompass distinct conformational changes upon state transitions, including rearrangements of local secondary structures in single-domain proteins, domain opening, domain rotations, and combinations of complex alterations in multi-domain proteins. As shown in Table 1, we used template-based modeling scores (TM-scores) and root-mean-square deviation (RMSD) to evaluate the conformation similarity (TM_2*states*_) and difference (RMSD_2*states*_) respectively. More details about the chosen proteins are elaborated in the Materials and Methods section. For consistency, we categorized these proteins into conformation 1 (C1) and conformation 2 (C2) in the representation of their two states, whose specific references and experimental structure information are listed in Table S1. We designated the apo-state as C1 and the holo-state as C2 and followed the definitions established in prior research for some membrane proteins, whose substrate-bounded or inward-facing state is usually defined as C2. 30–33

**Table 1.** AlphaFold2 prediction results for 20 selected two-state protein systems exhibiting significant conformational changes. Protein length, initial conformational similarity (TM_2*states*_), and difference (RMSD_2*states*_) between the two states are presented in columns two to four. The fifth column displays the number of sequences aligned using full MSA. The final two columns show the conformational differences between AlphaFold2 predictions and experimentally solved structures, represented as RMSD values relative to Conformation 1 (C1) or Conformation 2 (C2).

| Protein | Sequence Length | $TM_{2states}$ | $RMSD_{2states}$ (Å) | # MSA aligned sequences | $RMSD_{AF2}$ (Å) | |
| --- | --- | --- | --- | --- | --- | --- |
|  |  |  |  |  | C1 | C2 |
| AK | 214 | 0.69 | 6.95 | 17289 | 5.69 | 1.62 |
| LBP | 345 | 0.64 | 7.14 | 11060 | 6.82 | 0.64 |
| ALBP | 288 | 0.76 | 4.45 | 9168 | 3.77 | 1.46 |
| RBP | 271 | 0.76 | 4.06 | 9754 | 3.74 | 0.54 |
| GroEL | 524 | 0.64 | 12.9 | 20457 | 11.0 | 4.38 |
| GlnBP | 220 | 0.67 | 5.34 | 9143 | 5.15 | 0.76 |
| SPT10 | 480 | 0.82 | 4.40 | 14446 | 3.29 | 1.98 |
| DAHPS | 338 | 0.81 | 10.1 | 18906 | 9.69 | 1.32 |
| S100B | 83 | 0.55 | 5.93 | 8517 | 5.55 | 1.55 |
| CNBD | 142 | 0.74 | 6.09 | 10677 | 4.92 | 3.17 |
| NTRC | 124 | 0.64 | 4.55 | 17850 | 4.32 | 3.37 |
| AP4A | 165 | 0.80 | 4.61 | 13948 | 3.64 | 1.20 |
| 48G7g | 217 | 0.57 | 7.21 | 12360 | 6.30 | 2.13 |
| S100A1 | 93 | 0.50 | 6.39 | 10029 | 6.06 | 2.44 |
| T4L | 161 | 0.76 | 3.33 | 4078 | 0.92 | 2.50 |
| NTnC | 90 | 0.55 | 6.43 | 17122 | 3.90 | 4.14 |
| HSPD1 | 524 | 0.64 | 13.2 | 18296 | 3.01 | 12.7 |
| ZnT8 | 288 | 0.70 | 4.95 | 11863 | 2.96 | 3.19 |
| Mhp1 | 463 | 0.88 | 3.30 | 6057 | 2.30 | 3.08 |
| ASCT2 | 427 | 0.65 | 10.6 | 12605 | 3.57 | 7.74 |

We utilized ColabFold v1.5.2 (AlphaFold2 using MMseqs2) ^3,17^ to predict the structures and obtain corresponding distance maps (DMs) for the selected 20 proteins. Our approach involved a full-depth multiple sequence alignment (MSA) without recycling. We postulated that accurate protein structure prediction from coevolutionary information would encompass multiple functional states within the geometric conformational space. By skipping recycling, we preserved the original pair representations from the Evoformer and prevented updates that might bias the prediction towards a single, most probable conformation. Analysis of AlphaFold2’s default structure predictions revealed high precision in most models. However, these predictions often favored a single conformational state, aligning with recent findings. ^9,13^ This bias is evident in the last two columns of Table 1. For proteins exhibiting significant conformational changes between their two states, such as cyclic nucleotide-gated potassium channel (CNBD), transcriptional enhancer binding protein (NTRC), and regulatory domain of troponin C (NTnC), AF2’s predictions showed ambiguity. In these cases, the predicted structures deviated substantially from both known states. Given these observations, we sought to extract additional information from AF2’s output. We aimed to separate potential multistate information within its distance maps, enabling us to predict alternative conformations and potentially enhance the overall prediction accuracy of AlphaFold2.

We collected five default AF2 models for each protein, ranked from 001 to 005 based on their predicted local distance difference test (pLDDT) scores, following the protocols outlined in the Materials and Methods (Source of distance map). Our evaluation focused on the precision (Fig. 1A) and number abundance (Fig. 1B) of alternative conformational signals within the distance maps (DMs) across all 200 predicted models. For each of the 20 selected proteins, we analyzed five AF2 models in both prediction directions (from C1 to C2, and from C2 to C1). Our analysis revealed that the rank005 models overall represented the highest precision and abundance of signals associated with alternative conformations. Remarkably, even when the final predicted structures closely resemble a specific conformation, the rank005 DMs still contained significant information about alternative conformations. As illustrated in Fig. 1A, rank005 models exhibited the highest precision among rank001-005 models for 60% of the selected proteins. For example, Adenylate kinase (AK), D-allose binding protein (ALBP), Leucine binding protein (LBP), and Ribose binding protein (RBP) showed high-precision signals of alternative conformations in rank005 models, while other models showed almost no signals. Eight proteins, including NTnC, human mitochondrial chaperonin (HSPD1), S100 Calcium binding protein (S100B), DAHP synthase (DAHPS), Zinc transporter 8 (ZnT8), neutral amino acid transporter (ASCT2), S100 Metal binding protein (S100A1), and Ap4A hydrolase (AP4A) demonstrated higher precision in models other than rank005. However, when considering the ratio of signal numbers between other models and rank005 (Fig. 1B), except for DAHPS, seven of these proteins showed fewer or more sparsely distributed signals compared to rank005. This analysis suggests that a slight trade-off in precision may enhance the likelihood of capturing multiple conformations. Rank005 models performed well in capturing alternative conformation signals when considering both precision and quantity. Based on these results, we chose to use rank005 models for all tested proteins in subsequent analyses. Our results highlight the importance of examining lower-ranked models in AF2 predictions, as they may contain valuable information about alternative protein conformations that might be missed in higher-ranked models.

**Figure 1.**
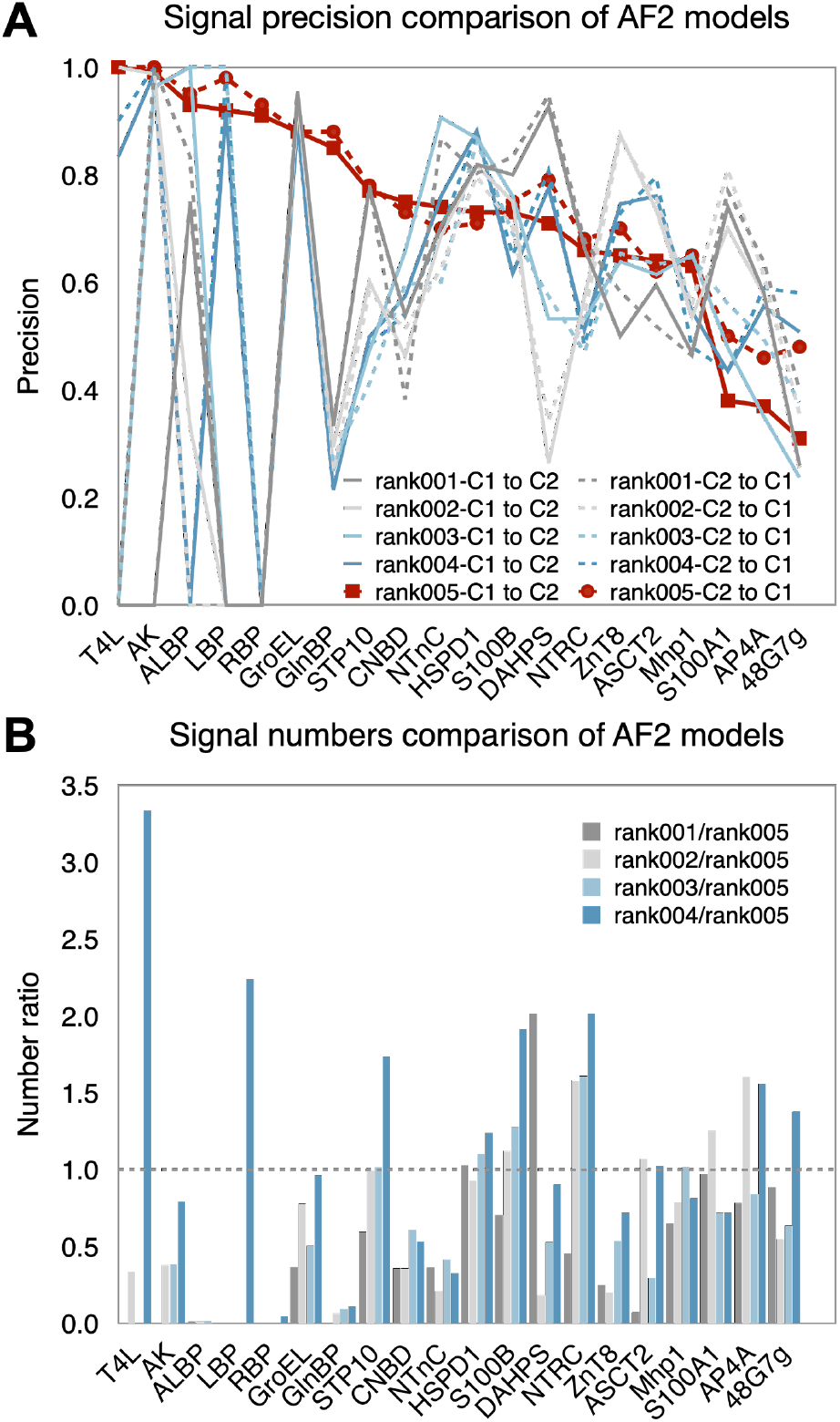
Evaluation of alternative conformation signals within the distance maps (DMs) of AF2 models. (A) Precision of alternative conformation signals for the default five models predicted by AF2. Overall, models rank005 demonstrate the best performance in multi-state prediction. Solid lines represent comparisons to conformation 1 (C1), while dotted lines indicate comparisons to conformation 2 (C2). (B) Number ratio of alternative conformational signals predicted by AF2 between rank001-004 and rank005 models. Most ratios fall below 1.0, suggesting that rank005 usually captures the most abundant signals of alternative conformations.

### 2.2 Extracting structural information of the alternative conformation based on a known structure and AF2 distance map

To utilize the alternative conformational information within AF2’s predictions, we developed a protocol for extracting multi-peak distribution signals. This method relies on distance maps predicted by AF2 and a known structure, as illustrated in Fig. 2. We use Adenylate kinase (AK) as an example to demonstrate this process. AF2’s default predicted structure for AK (shown in yellow) closely resembles the C2 (holo-state) conformation. However, we delved into AK’s predicted DM and uncovered numerous multi-peak distance-probability signals. These signals indicate distance variations between the same residue pairs, suggesting the presence of alternative conformations. Our approach assumes that the structure of one conformational state is already known, either through experimental determination or computational prediction. By comparing the multipeak signals from the AF2 prediction with this known structure, we can extract distinct distance distributions. These distributions then serve as guidance for predicting the second state.

**Figure 2.**
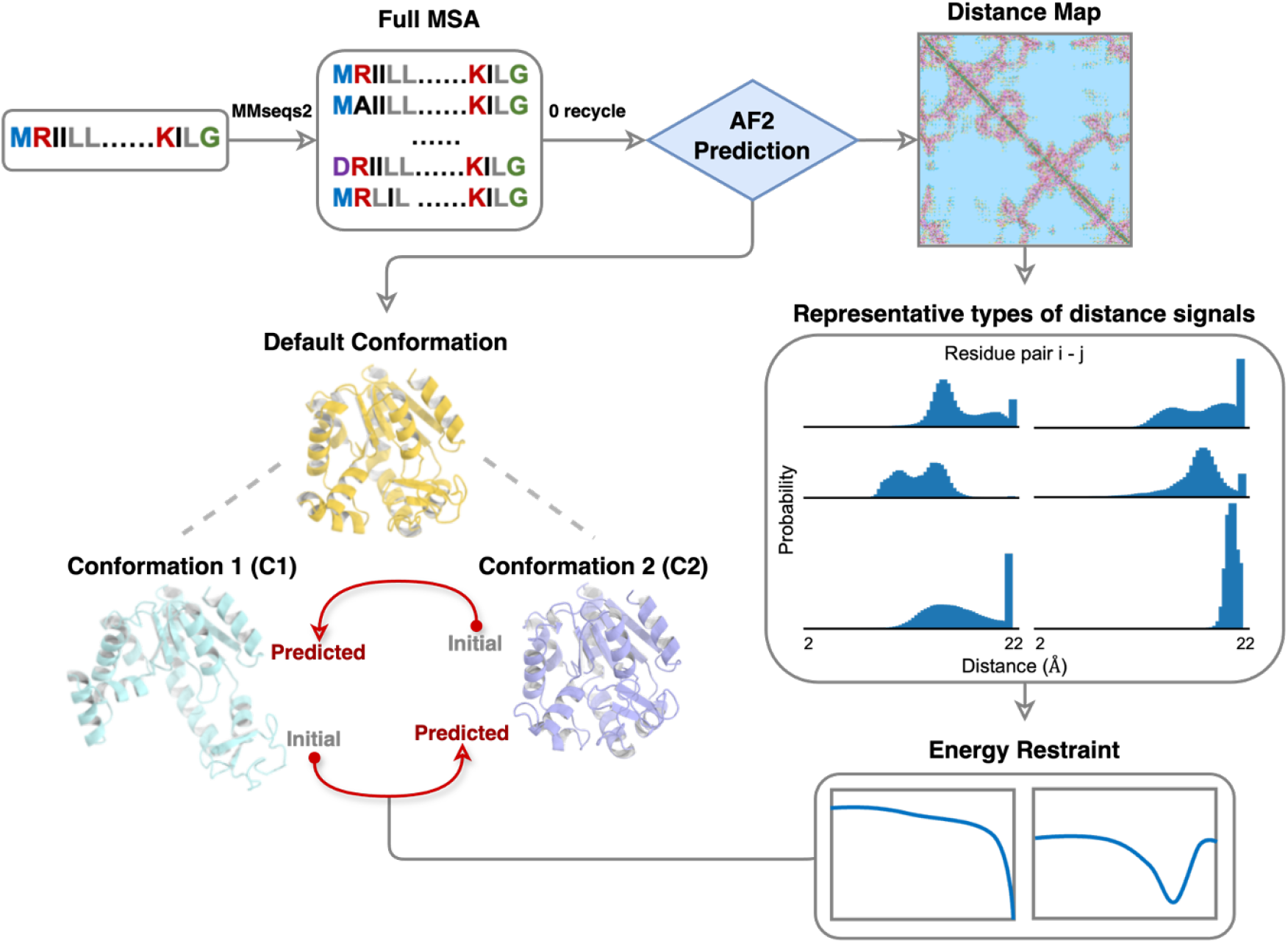
Workflow for predicting alternative conformations from a known structure and AF2 distance map (DM). AF2 was used to predict the default conformation and its DM, employing full-depth multiple sequence alignment (MSA) and zero recycle parameters. Starting from one state, we extracted and utilized multi-peak distance probability signals to predict the alternative state, enabling transitions from conformation 1 (C1) to conformation 2 (C2) and vice versa.

Our method for extracting alternative conformational information involves a detailed analysis of the distance probability distributions in AF2’s predictions. For residue pairs containing multipeak signals, we removed the distributions corresponding to the known state and rescaled the remaining distribution to a total probability of 1. Signals satisfying both conformational states were retained unchanged. These processed signals were combined to create a distance map for the alternative conformation (alt-DM), which was utilized in designing the restraint energy functions during the modeling process and guided our modeling of the second conformational state.

We categorized the distance-probability distributions into two main groups: 1) Multi-peak distributions, suggesting potential alternative distances between residue pairs and indicating conformational changes; 2) Single-peak distributions, representing stable structural elements across conformational states. We categorized six types of multi-peak distributions and thetypical singlepeak signals. Fig. 3 provides a comprehensive visualization of all these signal types. Detailed selection criteria are described in the Materials and Methods section. In brief, our signal extraction process focused on three aspects: 1) Fitting multi-peak distributions using a bivariate Gaussian Distribution 2) Employing the last bin’s data in distance maps (comprising 64 equal-width bins spanning from 2 to 22 Å) as an indicator of long-distance signals. 3) Incorporating single-peak signals that differ significantly from the known structure.

**Figure 3.**
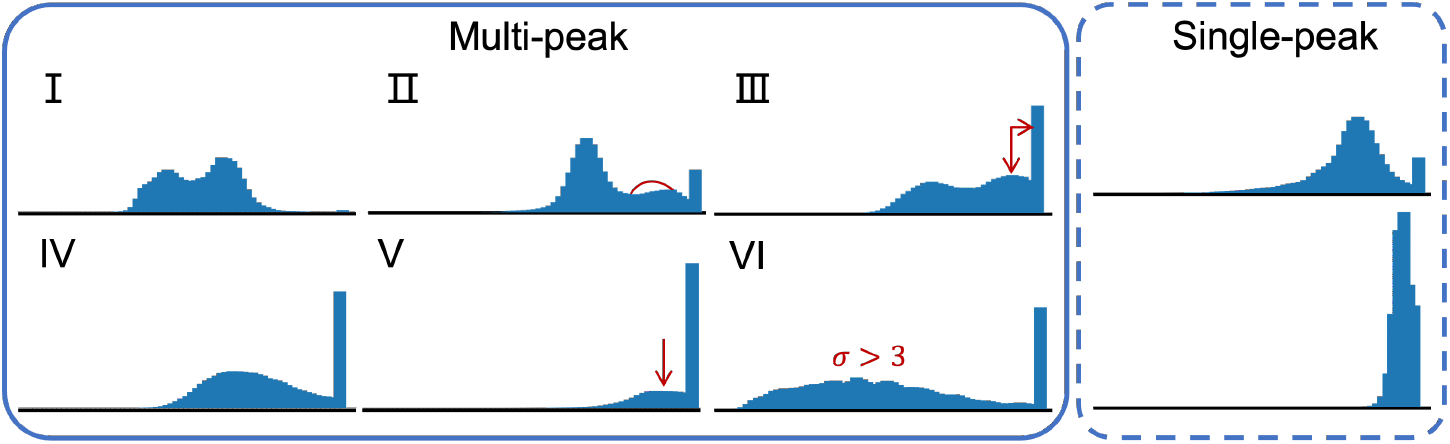
Categorization of distance-probability distributions. Multi-peak signals of six specific types and typical single-peak signals are illustrated. The corresponding selection criteria are described in the Materials and Methods section.

As evident from Table 2, AF2’s predictions for most of the studied proteins revealed an abundance of hidden signals of the alternative conformation. We analyzed the number of true positive and false positive of multi-peak signals (# Multi-peak), as well as the single-peak signals distinct from any known state (# Single-peak). For 13 out of 20 proteins, AF2 predictions with full MSA yielded hundreds of multi-peak signals,potentially valuable for modeling alternative conformations. An interesting outlier was T4 Lysozyme (T4L), which lacked multi-peak signals but contained four significant single-peak signals distinct from the known structure. Moreover, our signal selection protocol demonstrated remarkable precision in multiple instances, reaching nearly 1.0 for several multi-domain proteins. The average alternative conformation signal precision was 0.77 across the 20 proteins, effectively pinpointing the key residue pairs associated with the conformational differences between the two states.

**Table 2.**
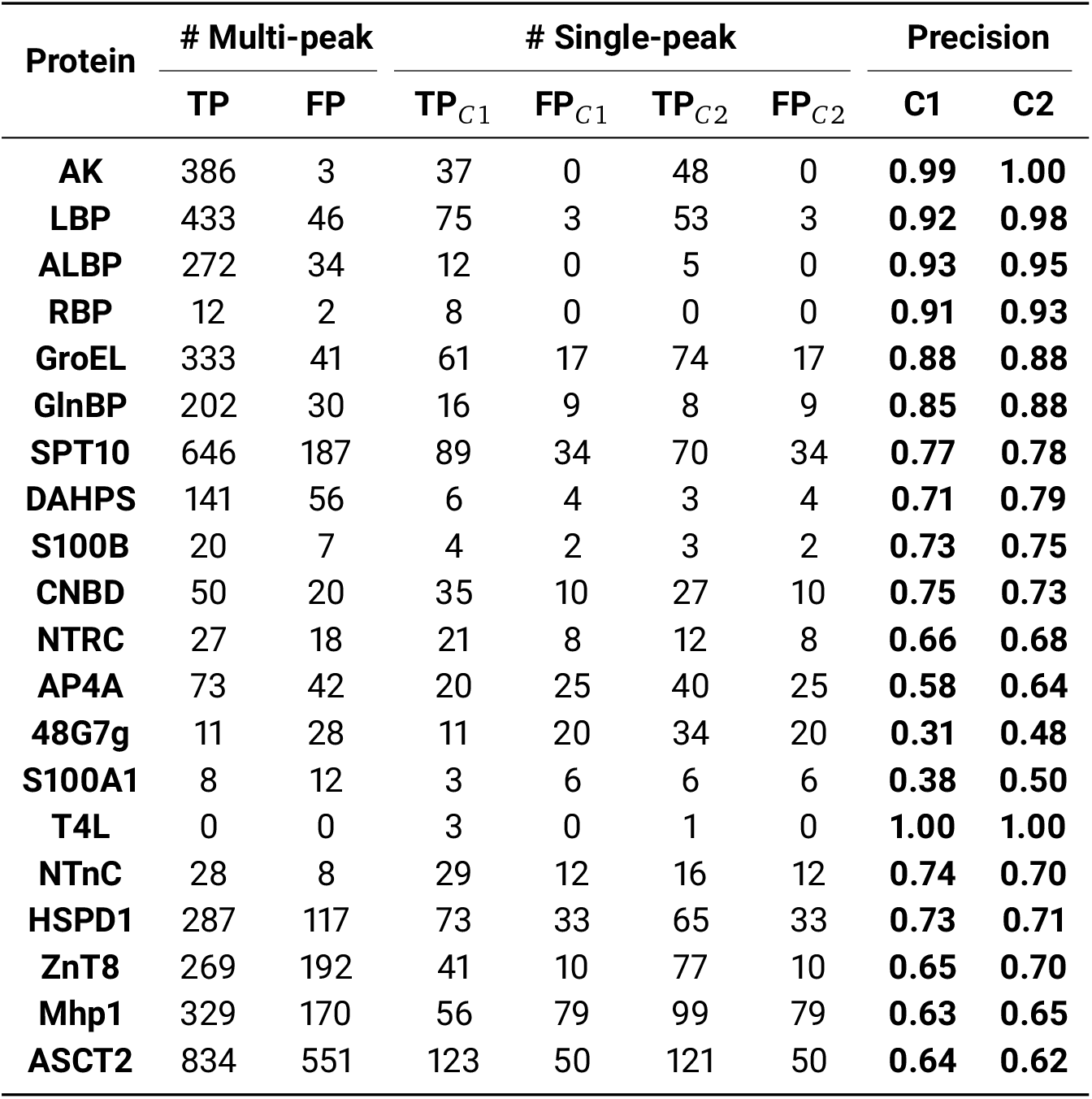
Analysis of alternative conformation signals from AF2 distance maps for 20 selected two-state proteins. The table quantifies signals that differ from the known state, categorized as multi-peak (# Multi-peak) and single-peak (# Single-peak) distributions. Precision scores, as defined in Eq. (3), are calculated based on known conformation 1 (C1 Precision) and known conformation 2 (C2 Precision). TP: true positive, FP: false positive.

| Protein | # Multi-peak |  | # Single-peak |  |  |  | Precision |  |
| --- | --- | --- | --- | --- | --- | --- | --- | --- |
|  | TP | FP | TP<sub>C1</sub> | FP<sub>C1</sub> | TP<sub>C2</sub> | FP<sub>C2</sub> | C1 | C2 |
| **AK** | 386 | 3 | 37 | 0 | 48 | 0 | **0.99** | **1.00** |
| **LBP** | 433 | 46 | 75 | 3 | 53 | 3 | **0.92** | **0.98** |
| **ALBP** | 272 | 34 | 12 | 0 | 5 | 0 | **0.93** | **0.95** |
| **RBP** | 12 | 2 | 8 | 0 | 0 | 0 | **0.91** | **0.93** |
| **GroEL** | 333 | 41 | 61 | 17 | 74 | 17 | **0.88** | **0.88** |
| **GlnBP** | 202 | 30 | 16 | 9 | 8 | 9 | **0.85** | **0.88** |
| **SPT10** | 646 | 187 | 89 | 34 | 70 | 34 | **0.77** | **0.78** |
| **DAHPS** | 141 | 56 | 6 | 4 | 3 | 4 | **0.71** | **0.79** |
| **S100B** | 20 | 7 | 4 | 2 | 3 | 2 | **0.73** | **0.75** |
| **CNBD** | 50 | 20 | 35 | 10 | 27 | 10 | **0.75** | **0.73** |
| **NTRC** | 27 | 18 | 21 | 8 | 12 | 8 | **0.66** | **0.68** |
| **AP4A** | 73 | 42 | 20 | 25 | 40 | 25 | **0.58** | **0.64** |
| **48G7g** | 11 | 28 | 11 | 20 | 34 | 20 | **0.31** | **0.48** |
| **S100A1** | 8 | 12 | 3 | 6 | 6 | 6 | **0.38** | **0.50** |
| **T4L** | 0 | 0 | 3 | 0 | 1 | 0 | **1.00** | **1.00** |
| **NTnC** | 28 | 8 | 29 | 12 | 16 | 12 | **0.74** | **0.70** |
| **HSPD1** | 287 | 117 | 73 | 33 | 65 | 33 | **0.73** | **0.71** |
| **ZnT8** | 269 | 192 | 41 | 10 | 77 | 10 | **0.65** | **0.70** |
| **Mhp1** | 329 | 170 | 56 | 79 | 99 | 79 | **0.63** | **0.65** |
| **ASCT2** | 834 | 551 | 123 | 50 | 121 | 50 | **0.64** | **0.62** |

### 2.3 Prediction of the alternative conformation with Rosetta

We further investigated whether the distance information identified above could aid in predicting the alternative state. With either C1 or C2 as a starting state, we constructed models for the alternative conformations of all 20 proteins in two directions (C1 to C2, or C2 to C1). This conformational exploration can be tackled through a physics-based optimization approach with the widely used Rosetta software suite.

#### 2.3.1 Convert DM into energy restraints for Rosetta modeling

The distance-probability distributions within DMs were converted into distance-based restraint energy functions and incorporated into Rosetta’s energy function during energy minimization and full-atom refinement stages. We constructed energy-distance functions (Fig. 4) based on the alternative distance maps (alt-DM) to guide the initial conformation towards the alternative state. Our sampling framework, which includes structure initialization, energy minimization, and full-atom refinement stages, resembles the protocol described in our recent work. ^29^

**Figure 4.**
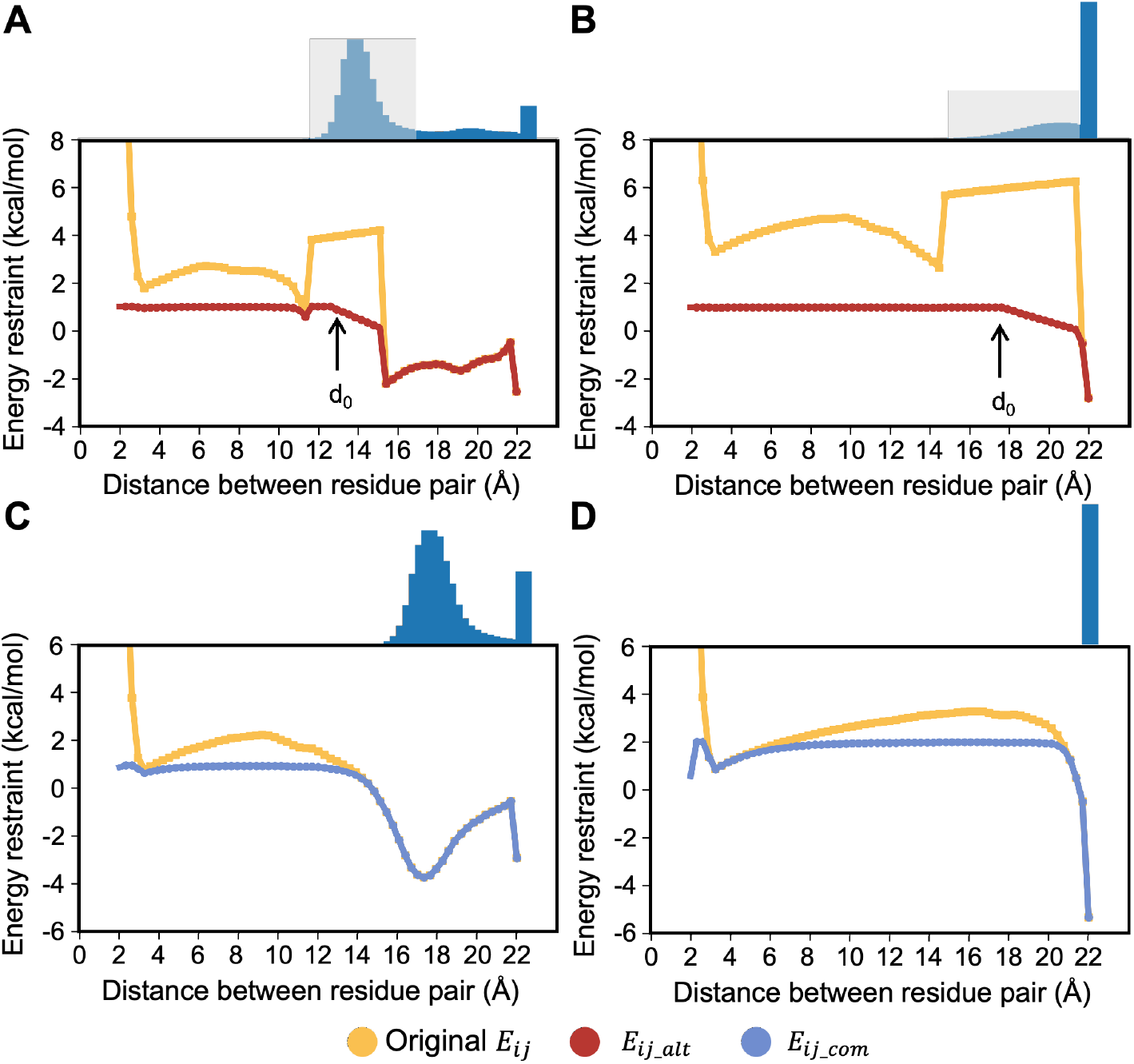
Restraint energy function for residue pairs with alternative (A-B) or common distributions between two states (C-D). Distance distributions from 2 to 22 Å correspond to blue signals at the top of each panel, with grey areas covering distance intervals specific to the initial state. The high energy barrier in the original distribution (yellow) is removed and smoothed in the designed energy function (red for *E*_*ij_alt*_ and blue for *E*_*ij_com*_). (A) Alternative state: predicted distance within the first 63 bins; initial distance in the known state (*d*_0_) = 12.7 Å. (B) Alternative state: predicted distance over 22 Å (last bin); *d*_0_ = 17.8 Å. (C) Common distribution: concentrated in the 16-18 Å range. (D) Common distribution: concentrated in the last bin (> 22 Å).

We distinguished between distance signals of alternative conformations (*E*_*i*j_*alt*_, Fig. 4A-B) and common signals shared by the two conformational states (*E*_*ij_com*_, Fig. 4C-D) in the restraints. As shown in Fig. 4, the energy barrier (yellow lines) associated with the known structure was flattened, facilitating changes in inter-residue distances during conformational transitions. For instance, the initial distance (*d*_0_) between the residue pair is 17.8 Å for Fig. 4B, while its predicted distance in the second state exceeds 22 Å, corresponding to the high probability distribution signal in the last bin.

A scaling parameter (*mag*) was used to balance the restraint strength between sparse alternative signals and common signals shared between two states. Residue pairs with precise distance predictions (lower comentropy *H*) were assigned higher weights, while signals with ambiguous distance predictions (higher comentropy *H*) were given less emphasis in the restraints. Signal density (*p*_*nl*_), defined as the ratio of alternative signals to sequence length, and the proportion of signals associated with the last bin among all alternative signals (*p*_*last*−*bin*_) were also considered in the parameter design. This approach aimed to effectively magnify restraints for alternative distributions while maintaining the overall structural rationality. For more detailed information, please refer to the ‘Structural modeling with Rosetta’ subsection in the Materials and Methods section.

#### 2.3.2 Assessment of predicted structures

After incorporating energy functions into the Rosetta modeling process, we sampled 150 models for each state of the 20 selected protein systems, using either C1 or C2 as the initial structure.

The sampling results, evaluated using Rosetta energy scores and TM-scores relative to each experimental structure, are displayed in Fig. 5 and Fig. 6. Initial structures are denoted by red stars, while sampled models are represented by circles colored according to their normalized energy scores. Models of inferior quality (*E*_*Rosetta*_>0) are marked with grey triangles. Dotted lines indicate the initial conformation similarities between the two states. The sampling distributions reveal that all proteins, except for ASCT2, tend to diverge from the initial red stars and approach the target conformational state (located at the opposite end of the diagonal).

**Figure 5.**
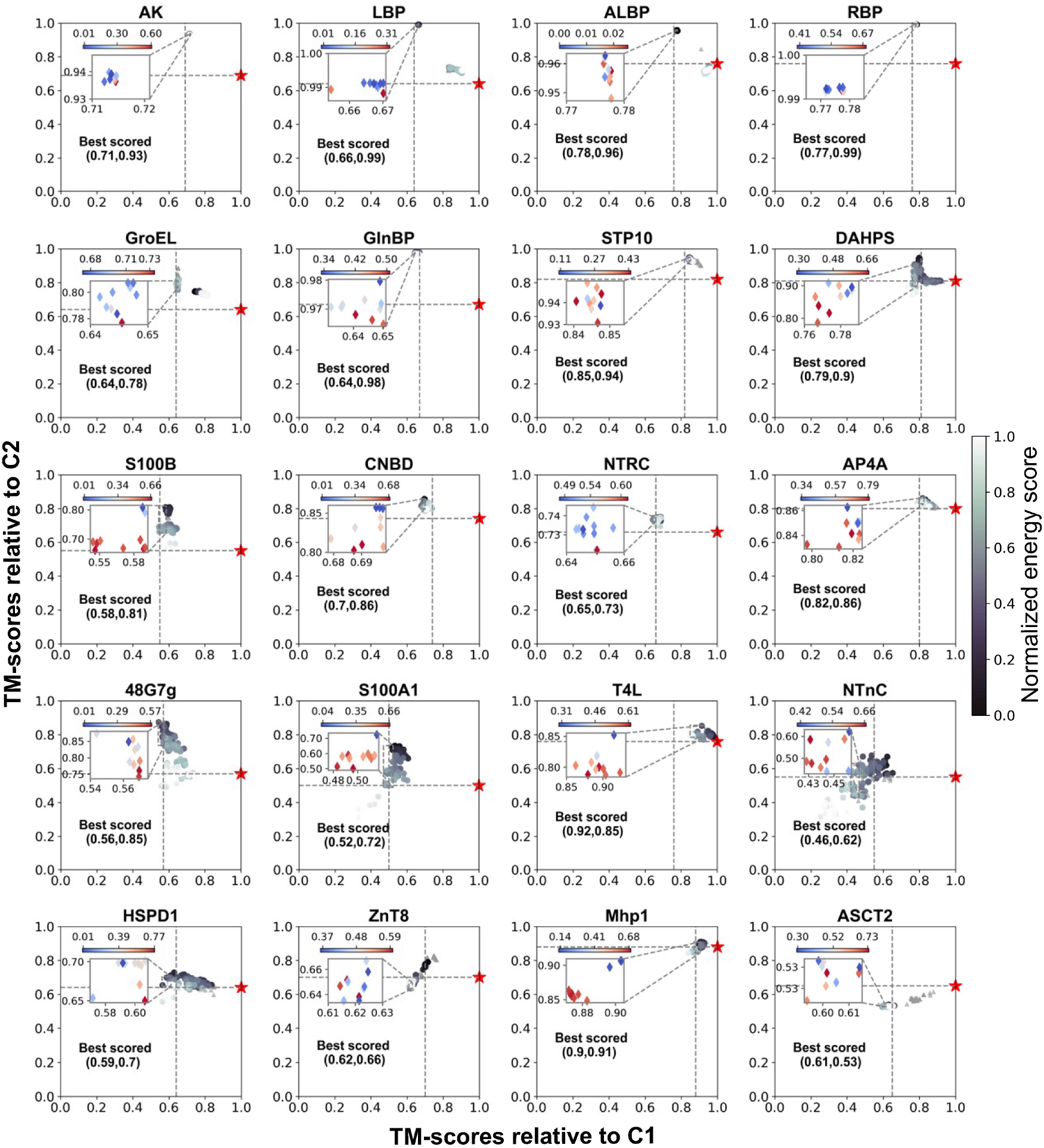
Rosetta sampling results for protein conformation prediction from C1 to C2. The initial structure (red star) and 150 models are shown, color-coded by normalized energy score (darker indicates lower energy). The dotted line represents the initial conformational similarity (TM_2*states*_) between states. Grey triangles denote structures not satisfying the designed restraints. Effective samples tend to shift across the diagonal. Inset: Top 10 models based on selection criteria, with normalized energy scores ranging from blue (lowest) to red (highest). The blue point indicates the best-scored model; its TM-scores relative to C1 and C2 are displayed in each panel.

**Figure 6.**
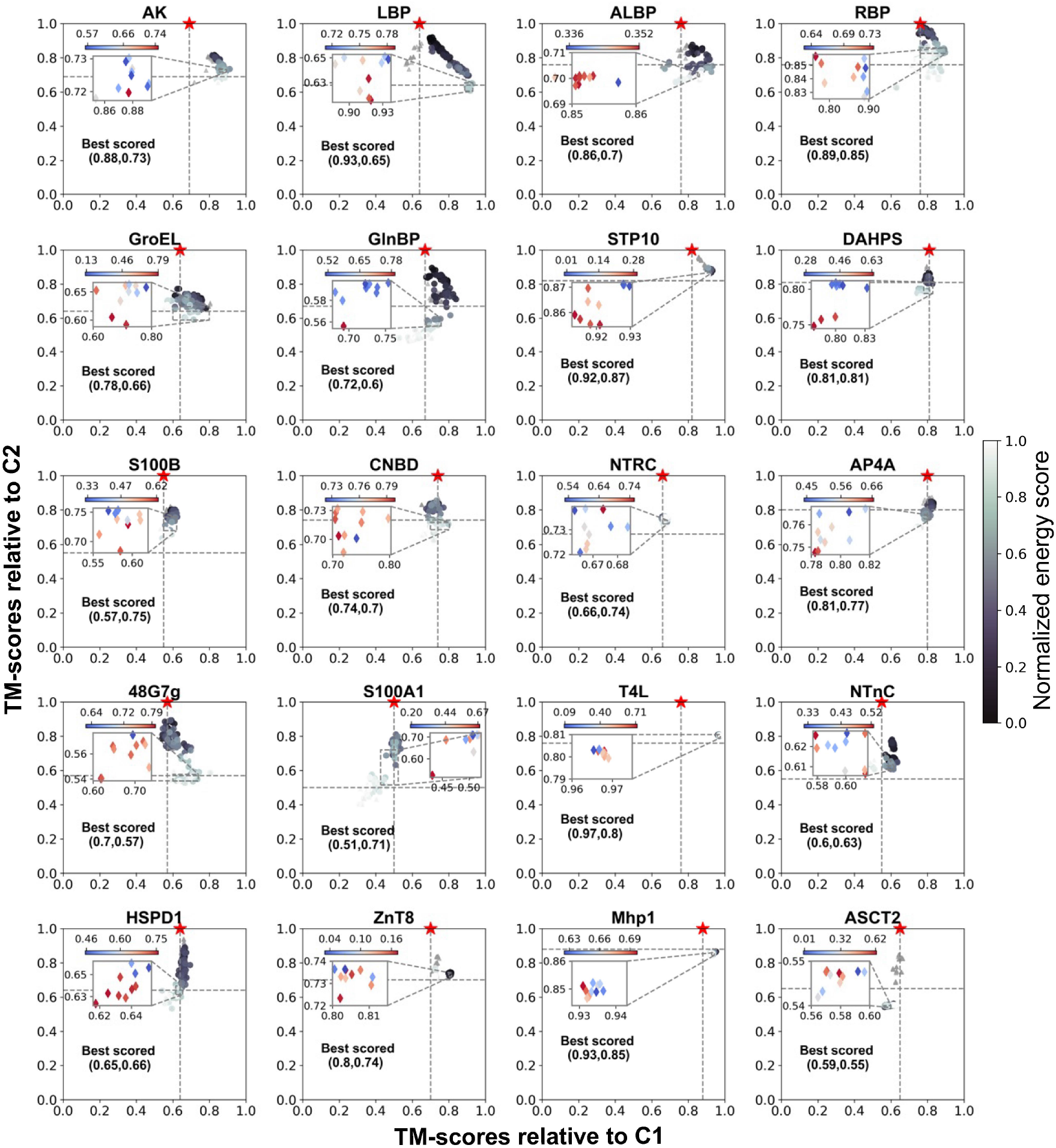
Rosetta sampling results for protein conformation prediction from C2 to C1. The initial structure (red star) and 150 models are shown, color-coded by normalized energy score (darker indicates lower energy). The dotted line represents the initial conformational similarity (TM_2*states*_) between states. Grey triangles denote structures not satisfying the designed restraints. Effective samples tend to shift across the diagonal. Inset: Top 10 models based on selection criteria, with normalized energy scores ranging from blue (lowest) to red (highest). The blue point indicates the best-scored model; its TM-scores relative to C1 and C2 are displayed in each panel.

We established criteria for selecting the best-scored models from a prediction standpoint (detailed in the Materials and Methods section). Priority was given to models exhibiting relatively lower energy scores and significant variations in TM-score compared to the initial structures. The top 10 models for each prediction, selected according to these criteria, are highlighted in the zoomed-in views, with colors ranging from blue to red. The blue point represents the best-scored model, and its TM-scores relative to C1 and C2 are labeled in each inset.

The quantitative prediction performance of our method is presented in Table 3, showcasing both the best-scored and best-sampled (best prediction among all samples) results for the two prediction directions. We considered a prediction successful if the TM-score relative to the target conformation exceeded the TM-score relative to the initial state. Successful predictions meeting this criterion are emphasized in bold font. When predicting from C1 to C2, the success rate was 85% (17/20) for the best-scored models and 90% (18/20) for the best-sampled models. In the opposite direction, from C2 to C1, the success rate was slightly lower at 70% (14/20) for both cases. Proteins with limited alternative signals (T4L) and significant conformational differences (*T M*_2*states*_ ≤ 0.55, S100B, S100A1, NTnC) posed greater challenges for predictions. The unsatisfactory TM-score performance of ASCT2 could be attributed to false positive signals in the last bins.

**Table 3.** Evaluation of predicted alternative states for 20 selected proteins. The initial conformational similarity between two states is represented by TM_2*states*_. Best-scored and best-sampled results are listed for both prediction directions. Superscripts denote the reference conformation for TM-score calculations. Successful predictions, defined as those closer to the target state than the initial structure, are highlighted in bold.

| Protein | $TM_{2states}$ | TM-score (C1 to C2) | | | | TM-score (C2 to C1) | | | |
| --- | --- | --- | --- | --- | --- | --- | --- | --- | --- |
|  |  | Best<sub>C1</sub><br>scored | Best<sub>C2</sub><br>scored | Best<sub>C1</sub><br>sampled | Best<sub>C2</sub><br>sampled | Best<sub>C1</sub><br>scored | Best<sub>C2</sub><br>scored | Best<sub>C1</sub><br>sampled | Best<sub>C2</sub><br>sampled |
| AK | **0.69** | 0.71 | **0.93** | 0.71 | **0.94** | **0.88** | 0.73 | **0.91** | 0.73 |
| LBP | **0.64** | 0.66 | **0.99** | 0.66 | **0.99** | **0.93** | 0.65 | **0.93** | 0.65 |
| ALBP | **0.76** | 0.78 | **0.96** | 0.78 | **0.96** | **0.86** | 0.70 | **0.93** | 0.75 |
| RBP | **0.76** | 0.77 | **0.99** | 0.77 | **0.99** | **0.89** | 0.85 | **0.90** | 0.83 |
| GroEL | **0.64** | 0.64 | **0.78** | 0.65 | **0.88** | **0.78** | 0.66 | **0.80** | 0.69 |
| GlnBP | **0.67** | 0.64 | **0.98** | 0.65 | **0.98** | **0.72** | 0.60 | **0.84** | 0.71 |
| SPT10 | **0.82** | 0.85 | **0.94** | 0.84 | **0.95** | **0.92** | 0.87 | **0.93** | 0.88 |
| DAHPS | **0.81** | 0.79 | **0.90** | 0.80 | **0.95** | **0.81** | 0.81 | **0.83** | 0.80 |
| S100B | **0.55** | 0.58 | **0.81** | 0.60 | **0.81** | 0.57 | 0.75 | 0.62 | 0.76 |
| CNBD | **0.74** | 0.70 | **0.86** | 0.70 | **0.86** | **0.74** | 0.70 | **0.80** | 0.72 |
| NTRC | **0.66** | 0.65 | **0.73** | 0.68 | **0.75** | 0.66 | 0.74 | 0.69 | 0.75 |
| AP4A | **0.80** | 0.82 | **0.86** | 0.82 | **0.86** | **0.81** | 0.77 | **0.83** | 0.83 |
| 48G7g | **0.57** | 0.56 | **0.85** | 0.58 | **0.87** | **0.70** | 0.57 | **0.80** | 0.59 |
| S100A1 | **0.50** | 0.52 | **0.72** | 0.54 | **0.74** | 0.51 | 0.71 | 0.54 | 0.73 |
| T4L | **0.76** | 0.92 | 0.85 | 0.92 | 0.85 | **0.97** | 0.80 | **0.97** | 0.80 |
| NTnC | **0.55** | 0.46 | **0.62** | 0.54 | **0.68** | 0.60 | 0.63 | 0.64 | 0.65 |
| HSPD1 | **0.64** | 0.59 | **0.70** | 0.63 | **0.73** | 0.65 | 0.66 | 0.70 | 0.79 |
| ZnT8 | **0.70** | 0.62 | 0.66 | 0.75 | **0.83** | **0.80** | 0.74 | **0.81** | 0.74 |
| Mhp1 | **0.88** | 0.90 | **0.91** | 0.91 | **0.91** | **0.93** | 0.85 | **0.95** | 0.87 |
| ASCT2 | **0.65** | 0.61 | 0.53 | 0.88 | 0.62 | 0.59 | 0.55 | 0.66 | 0.82 |

The prediction capability of our protocol for alternative conformations was demonstrated across proteins of various sizes and conformational changes, as illustrated in Fig. 7 and Fig. 8. Fig. 7 provides structural visualizations, showing AF2’s predicted models in yellow, representative experimental C1 structures in grey, and C2 structures in blue. The predicted C1 (green) and C2 (purple) structures are aligned to their target states, respectively, visually demonstrating the prediction precision. For proteins exhibiting relatively simple changes, such as domains “opening” and “closing” through hinge-bending (LBP, ALBP, RBP), we achieved an average precision of 0.98 for C2 and 0.92 for C1. In these cases, AF2 provided remarkably accurate predictions closely resembling C2, while our successful modeling of C1 was highly valuable, predicting the alternative conformation not directly provided by AF2. More interestingly, for proteins involving domain rotation (DAHPS, Immunoglobin 48G7 germline Fab—48G7g) and complex conformational changes (AK, 60 kDa chaperonin—GroEL), the effectiveness of the designed energy restraints can be inferred from valid predictions in both directions as well. For example, the initial TM-score between the two states of 48G7g is 0.57, indicating a major conformational difference through domain rotations (Fig. 7). Our predicted C1 and C2 structures achieved TM-scores of 0.80 and 0.87 relative to the experimental structures, respectively, correctly capturing the rotation tendency between domains while maintaining accurate structure prediction within single domains. However, in certain challenging cases (e.g., ASCT2, S100A1), our method proved less efficient in rearranging helices due to limited precision and sparse signals from the alt-DM analysis.

**Figure 7.**
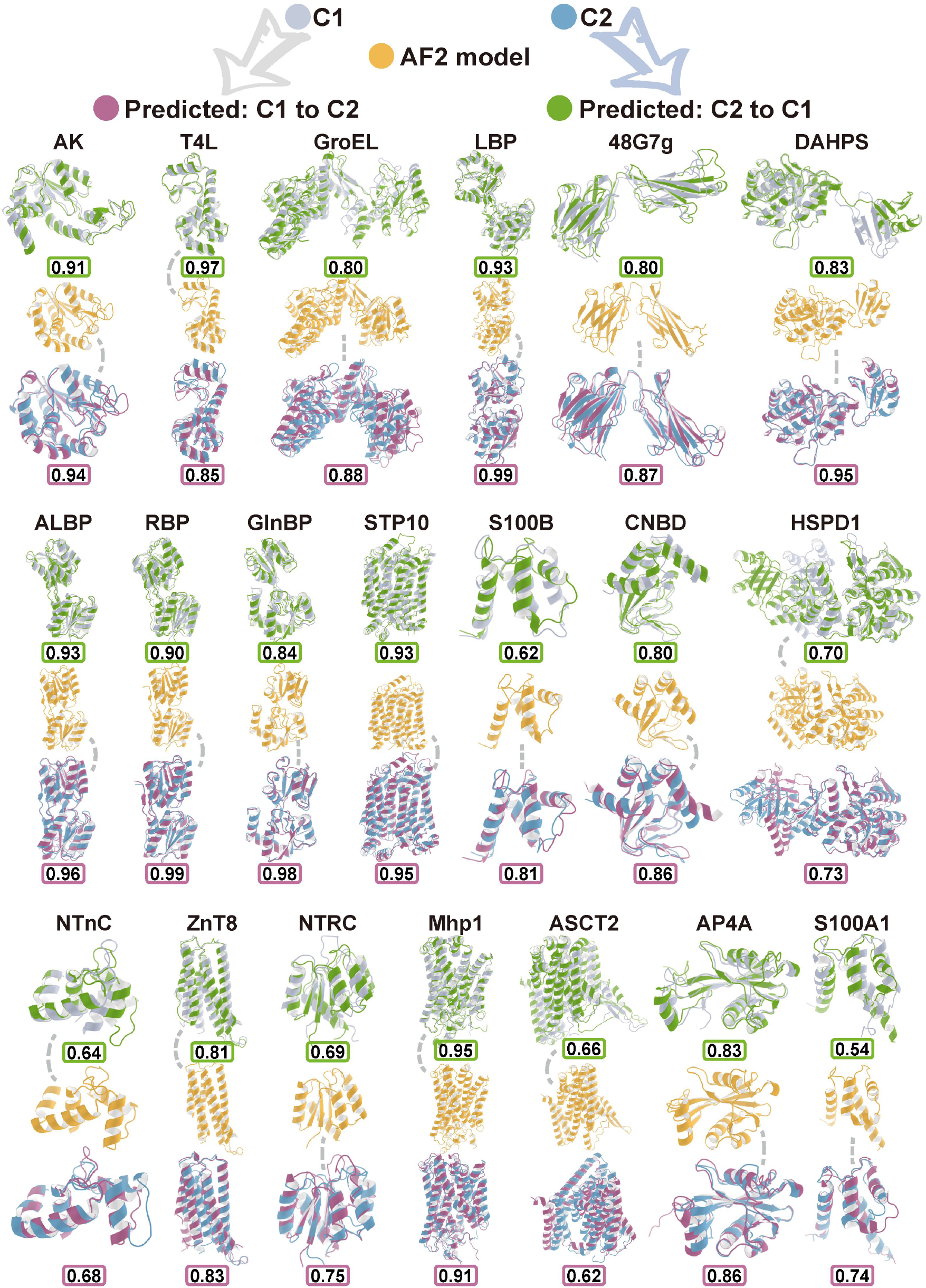
Predictions for the 20 selected proteins with two conformational states. Initial experimentally solved structures (C1: grey; C2: blue) are aligned with best-sampled structures (predicted C1: green; predicted C2: purple). The default AF2 models (yellow) are shown in the middle rows, connected by dotted lines to conformations with higher similarity. TM-scores for predictions are shown in color-coded boxes (green for C1, purple for C2). Nine out of twenty proteins achieved TM-scores above 0.90.

**Figure 8.**
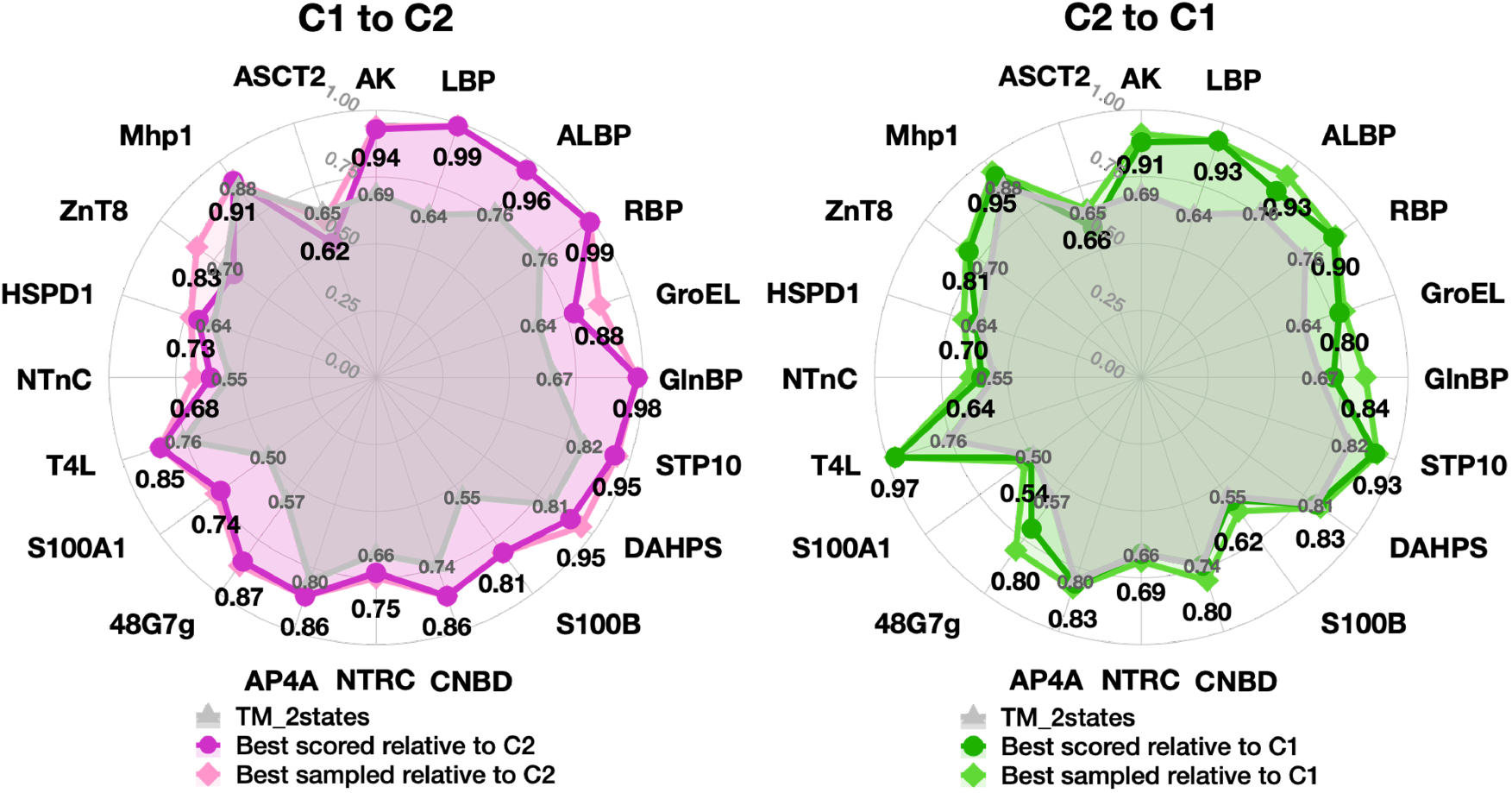
Radar maps of TM-scores for the 20 proteins in bidirectional predictions. C1 and C2 represent Conformations 1 and 2, respectively. The grey region indicates the initial conformational similarity (TM_2*states*_) between the states. Colored shapes show the TM-scores between the predicted structures and the target state, with darker-colored boarders representing the best-scored model and lighter-colored boarders the best-sampled model.

Fig. 8 presents a radar map of the TM-score results for the 20 proteins. The grey area indicates the initial conformation similarities between the two states, while the best-scored (darker colored lines) and best-sampled (lighter colored lines) results in both prediction directions cover larger areas, demonstrating that the modeling successfully steered the structure from one state toward the other. For the prediction from C1 to C2, our predicted structures became closer to the target C2 conformation in 19 out of 20 systems (ASCT2 being the exception), achieving high precision (TM-score > 0.90) in eight protein systems. Even in the more challenging direction (C2 to C1), where most target conformations deviate significantly from AF2’s predictions, we successfully predicted conformations approaching C1, with seven proteins reaching a TM-score over 0.90. The average TM-scores for predictions in the C1 to C2 and C2 to C1 directions were 0.86 and 0.80, respectively, across the 20 protein systems. These results demonstrate the effectiveness of our protocol in modeling two-state proteins with various types of significant conformational changes.

We conducted two control experiments to assess the effectiveness of our method. The first control experiment (CTRL1) used the entire AF2 distance maps for modeling, without our signal selection protocol. The second control experiment (CTRL2) omitted the scaling parameter ‘mag’ in the energy restraint function, treating all signals with equal weights. Fig. S1 compares the sampling results of these control experiments with our method. Dotted lines depict the TM-score relative to the initial state in each prediction direction, while solid lines represent the TM-score relative to the target state. Our sampling results exhibited higher precision compared to both control experiments in 18 out of 20 proteins for the C1 to C2 direction and in all 20 proteins for the C2 to C1 direction (as shown by the more peripheral purple and green solid lines). Notably, in the control experiments, the dotted lines were more peripheral, indicating that the predicted conformations were biased towards the initial structures rather than transitioning to the alternative state. This observation supports the effectiveness of our prediction method, which uses selected signals of the alternative conformation and balances the energy function with higher weights for these signals. Detailed results and sampling plots can be found in Tables S2-S3 and Figures S2-S5.

## 3 Discussion

In recent years, deep learning-based protein structure prediction has exhibited remarkable accuracy, particularly for simple proteins. ^3,4^ A limitation of most current methods is their focus on predicting a specific structure, neglecting the existence of multiple-state conformations, while structural heterogeneity underpins protein functions. ^34^ However, this limitation does not indicate that deep-learning methods are incapable of predicting multiple conformations. We believe in their vast potential for this task. The prediction of alternative states is gaining increasing attention from various research perspectives. ^13,18,21,22,24,29,35,36^ In this work, our results revealed that AlphFold2 predictions inherently contain latent information about the alternative structural states for the proteins with multiple functional states.

With this hidden information extracted and combined with Rosetta, we successfully predicted alternative conformations for 19 two-state proteins starting from either C1 or C2. Our method’s effectiveness is evident when the target conformations differ significantly from those predicted by AF2, demonstrating our protocol’s ability to identify and utilize distance distributions of the alternative conformations for structure modeling. However, one crucial factor influencing prediction quality is the initial DM predicted by AF2. As we rely on AF2 models generated without recycles, not all test proteins achieved high precision. Ambiguous conformations between two functional states predicted by AF2 can sometimes introduce numerous false signals. For example, the RMSDs of zinc transporter ZnT8 predicted by AF2 relative to its two states are 2.96 and 3.19 Å, which do not correspond to either state. The precision of ZnT8’s alternative signal prediction from two directions both fall below 0.70, further highlighting the challenge of distinguishing between functional conformations in such cases. The presence of false positive distributions on structurally important residue pairs, especially those separated by long distances exceeding 22 Å, can compromise the precision of the conformation sampling process. To address limited prediction precision in certain proteins, we anticipate that deep-learning models other than Rosetta, such as Distance-AF ^37^ or Chroma, ^38^ may enhance the utilization of extracted alternative information from the DMs.

There were other studies that investigated some of the protein systems in our test set. ^22,24^ We conducted a comparison between the best results reported in those papers and the predictions of our work. As shown in Table 4, proteins AK and RBP were tested using two other prediction methods. We compared the prediction accuracy with the reported RMSD or TM-score. C1 of both proteins corresponds to the apo state, which is a relatively difficult prediction target. Our study achieved a prediction accuracy for AK’s C1 with RMSD=1.84 Å and TM-score=0.91, surpassing the prediction accuracy reported by DiG and SPEACH_AF. ^22,24^ For the simpler C2 prediction, our results achieved comparable prediction accuracy to these two studies. For example, both methods achieved a TM-score of 0.99 for RBP’s C2, demonstrating the capability of these prediction methods. However, the prediction performance for several transmembrane proteins in our study was a bit inferior. The average prediction accuracy for Sugar symporter (STP10), ZnT8, and ASCT2 in other AF2-based subsampling methods was above 0.90 (TM-score). ^18,22^ Although our method achieved an accurate prediction with TM-score >0.90 for STP10, the accuracies for ZnT8 and ASCT2 were less satisfactory, only 0.81 and 0.62, respectively (Table 4). For ZnT8 and ASCT2, the modeling was affected by false positive signals for the alternative conformations (Table 2), indicating that our method still had room for improvement.

**Table 4.** Comparison of prediction performance on the same representative proteins with other deep-learning-based methods.

| Method | Protein | RMSD/TM-score<br>in papers |  | RMSD/TM-score<br>in this work |  |
| --- | --- | --- | --- | --- | --- |
|  |  | Relative to<br>C1 | Relative to<br>C2 | Relative to<br>C1 | Relative to<br>C2 |
| DiG<sup>24</sup> 2023 | AK | <3 Å | <1 Å | **1.84 Å** | 1.51 Å |
| SPEACH_AF<sup>22</sup> 2022 | AK | 0.89 | 0.98 | **0.91** | 0.94 |
|  | RBP | 0.98 | 0.99 | 0.90 | 0.99 |
|  | ZnT8 | 0.95 | 0.92 | 0.83 | 0.81 |
|  | ASCT2 | 0.96 | 0.92 | 0.66 | 0.62 |
|  | STP10 | 0.98 | 0.98 | 0.93 | 0.95 |

To validate our prediction protocol, we conducted a control experiment using two extremostable proteins. ^39^ As shown in Fig. S6, there was no signal for protein Hsp31 (PDB: 1izy) and only sparse signals on the terminal loop for protein FGF1 (PDB: 1axm), which were insufficient to suggest a distinct second state. These findings support the robustness of our protocol, demonstrating that it does not generate unrealistic conformations for proteins with a single stable state.

However, our protocol inherently cannot deal with the challenging cases when proteins possess more than two states. Encouragingly, MD-based enhanced sampling procedures, ^40–43^ smarter MSA subsampling strategies, ^21^ and deep learning-based generative models ^24^ may provide more comprehensive pictures for multi-state structures. Therefore, we are optimistic that the rapid development of the field and probably the combination of the above methods can solve the problem in the near future.

In summary, this study explored the hidden multiple-state structural information within AF2’s predictions without shallowing MSA and established an approach to incorporate alternative conformation signals derived from deep learning predictions into physics-based modeling. We tested a wide range of structural variations and protein sizes, which demonstrated the effectiveness of our protocol. This framework can serve as a rapid hypothesis-testing tool for exploring the likely alternative conformation of a protein structure or predicting two most likely structures from a given sequence. Experimental techniques such as protein conjugation, NMR, and FRET can also be combined with our protocol to unravel the dynamics and functions of multistate proteins.

## 4 Materials and Methods

### 4.1 Overall protocol

1. As shown in Fig. 2, we extracted distance maps (DMs) from AlphaFold2’s predictions. These DMs served as the foundation for identifying residue pair distances associated with alternative conformations.
2. Starting from either state, we calculated its corresponding distance map and compared it to the AlphaFold2 prediction. Signals indicating alternative conformations were recorded in the alt-DM (Distance Map for alternative conformation).
3. We used the alt-DM for further structure modeling and designed the distance-based energy restraint function that accounted for both alternative or common conformational signals.
4. We generated a series of structural models using Rosetta, incorporating the restraint energy function. These models underwent evaluation to give the best model.

### 4.2 Protein systems studied in this work

Our study utilized a dataset of 20 two-state proteins, selected from databases curated by Krebs et al. ^44–46^ and Tadeo et al. ^13^ This diverse set of proteins exhibits various types of conformational changes (details in Table S1), which we categorized as follows:

- Hinge-bending: These proteins display straightforward ‘open’ and ‘closed’ domain movements. Examples include Leucine binding protein (LBP), Glutamine-binding protein (GlnBP), Ribose binding protein (RBP), D-allose binding protein (ALBP).
- Inter-domain rotations: This category includes proteins such as DAHP synthase (DAHPS), Immunoglobin 48G7 germline Fab (48G7g), which exhibit rotational movements between domains.
- Single-domain local rearrangements: Several proteins in our dataset show localized changes in helices and loops within a single domain. Examples are S100 Calcium-binding protein (S100B), Nucleotide-activated K^+^ channel binding domain (CNBD), Receiver domain of transcriptional enhancer binding protein (NTRC), Ap4A hydrolase (AP4A), S100 Metal-binding protein (S100A1), T4 Lysozyme (T4L), Regulatory domain of troponin C (NTnC). Also, there are some membrane proteins including Sugar symporter (STP10), Zinc transporter 8 (ZnT8), Hydantoin transport protein (Mhp1), and Neutral amino acid transporter (ASCT2).
- Complex multi-domain rearrangements: Some proteins in our dataset, such as Adenylate kinase (AK), 60kDa chaperonin (GroEL), and Human mitochondrial chaperonin (HSPD1), exhibit a combination of different conformational changes across multiple domains.

Detailed information about all structures, downloaded from the wwPDB, ^47^ is summarized in Table S1.

### 4.3 Source of distance maps

We used ColabFold v1.5.2: AlphaFold2 using MMseqs2^17,48^ for structure predictions of all systems. To optimize accuracy and explore potential multi-state conformations, we implemented the following parameter choices: (1) We chose full MSA depth as *max_msa*=512:1024 in advanced settings; (2) We disabled recycling (*num_recycles*=0) to focus on initial predictions; (3) No templates were used in our predictions; (4) We used a single seed (*num_seeds*=1), resulting in five models per prediction process; (5) We selected the ‘save_all’ option, with particular focus on the ‘.pickle’ file for distance map extraction. All other parameters adhered to ColabFold’s default settings.

AlphaFold’s algorithm converts pair representations into 64 distance bins, ^48^ assigning probabilities using a *softmax* function. These bins span distances from 2 to 22 Å in equal increments, with the final bin encompassing all distances beyond 22 Å. To calculate the probability of distances between residue pairs, we processed the [‘distogram’][‘logits’] data from the ‘.pickle’ file using the softmax equation Eq. (1). This calculation was performed for *C*_*β*_ atoms of all residue pairs, except for glycine, where we used the *C*_*α*_ atom due to the absence of a *C*_*β*_.

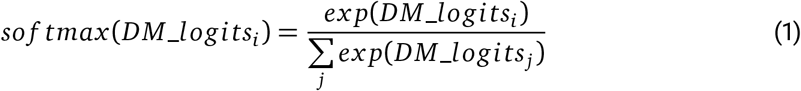

### 4.4 Extracting information on alternative conformations from DM analysis

To identify potential alternative conformations in protein structure predictions, we developed a systematic approach for analyzing the complex features exhibited in the DMs. Our method focuses on detecting and classifying multi-peak signals in the predicted distance-probability distributions. The classification criteria and corresponding signal types are illustrated in Fig. 3. Our analysis procedure consists of three major steps:

1. Initial peak detection and classification: For each residue pair *i*− *j*, we examine the first 63 distance bins, reserving the last bin for long-distance signal analysis. We identify local maxima and select the top two (*max* 1, *max* 2). These peaks are checked based on their relative probabilities and separation: the probability sum of two distinct peaks should exceed 0.1; the probability of two maxima should satisfy *p*_*max*2_ *>* 0.1*p*_*max*1_; closely spaced peaks (*p*_*max*1_ − *p*_*max*2_ *<* 0.05*p*_*max*1_) requiring consideration of a third maximum.
2. Bivariate Gaussian Distribution fitting: For potential multi-peak signals, we apply a bivariate Gaussian distribution fit (Eq. (2)) with the following initial parameters. *µ*1, *µ*2 are corresponding distances of two maxima, the initial amplitude *a*1 = *a*2 = 0.1, and standard deviations *σ*1 = *σ*2 = 1.

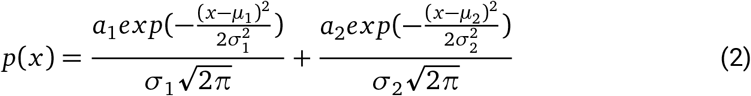 According to the fitting results, we classify the signals into the following categories:
  - Class I: Peaks with *σ* < 3, ensuring distinct distributions.
  - Class II: Secondary peak is weak but significantly above the inter-peak saddle point ((*p*_*max* 2_ − *p*_*saddle*_)*/p*_*max* 2_ *>* 0.05).
  - Class III: Peak is well-defined and falls within appropriate distance ranges (*µ* + *σ <* 21.6875 Å).
3. Long-distance signal analysis: We assess the probability of the last distance bin (*p*_*last*_) to identify significant long-distance signals. A notable but not dominant long-distance signal is indicated when *p*_*last*_ is larger than any fitted amplitude and falls between 0.1 and 0.8.

For distributions with a single main peak, we apply a univariate Gaussian fit and similar classification criteria. Multi-peak cases involving long-distance signals are classified as Class IV to VI, depending on the fitting results:

- Class IV: Single peak from one maximum in step 1.
- Class V: Only one peak fitted by Eq. (2) in step 2.
- Class VI: Peaks fitted in step 2 with *σ* > 3 are combined as one single peak.

Following this signal classification criteria, we further analyze the signals to identify and characterize alternative conformations:

a. Peak analysis: For each residue pair, we determine the number of signal peaks (*n*_*peak*_) and their corresponding breadths. Residue pairs with *n*_*peak*_ *>* 1 are considered potential indicators of alternative conformations.
b. Comparison with known structures: We compare predicted DMs with known structures to validate the signals.

- For multi-peak signals (*n*_*peak*_ *>* 1), we remove the peak corresponding to the distance in the known structure and retain the remaining signals in alt-DM.
- For single-peak signals (*n*_*peak*_ = 1), we compare the peak breadth with the known distance. If a discrepancy is found, we include this signal in our further modeling.
c. Signal classification and precision calculation: We classify signals effective in predicting the alternative state into two categories, true positive multi-peak signals (*T P*_*m*_) and true positive single-peak signals (*T P*_*s*_). Both categories contribute to our final precision calculation (Eq. (3)), where *C*_*i*_ represents conformational state *i*.

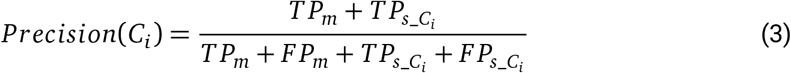

### 4.5 Model building and assessment

#### 4.5.1 Structural modeling with Rosetta

Starting with one known protein structure, e.g. C1, we used Rosetta to build plausible models of the alternative conformation with restraints derived from the above DM signals. Three functional modules of PyRosetta, ^49^ pose initialization, minimization, and full-atom refinement were employed in this process.

We initialized the pose using the “centroid” switch type. To accommodate our distance-energy function, we mutated glycine residues to alanine, ensuring all residues had *C*_*β*_ atoms for distance calculations.

We developed piece-wise energy-distance functions to represent restraints from alt-DM signals, adapting the probability-to-energy transition method from trRosettaX. ^6^ Our approach differentiates between residue pairs with alternative conformation signals and those with common distributions between two states:

1. For alternative conformation signals: As shown in Fig. 4A-B, we flattened energy barriers at locations corresponding to the initial distance peak using a hyperbolic tangent function (*E*_*i*j_*temp*_, Eq. (4)). We then applied gradient descent toward the new distribution (*E*_*ij_alt*_, Eq. (5)) based on the initial distance between residue pairs (*d*_*ij*_).

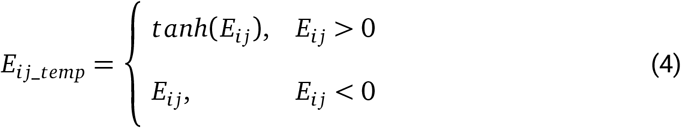

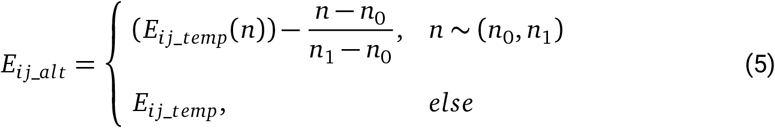

where

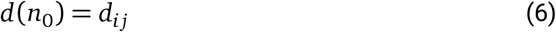

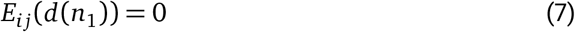

2. For common distributions between two states: As shown in Fig. 4C-D, we slightly flattened energy barriers above a platform value (Eq. (9)), where *Mo* (Eq. (8)) is the statistical mode of positive energy values.

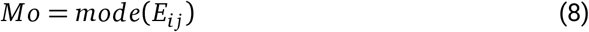

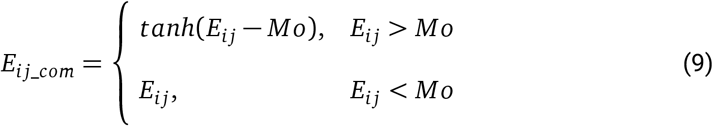

To enhance the influence of alternative signals, we introduced a scaling parameter ‘mag’ (Eq. (10)) that considers *p*_*nl*_ (the ratio of alternative signals to protein length, Eq. (11)), *p*_*last*−*bin*_ (the proportion of long-distance signals, Eq. (12)), and *H*_*norm*_ (normalized comentropy, Eq. (14)). Among these equations, *N*_*alt*_ is the number of signals connected to alternative conformations; *N*_*last*−*bin*_ is the number of signals in the last bin, which represents a large distance separation exceeding 22Å; *H*(*p*) is the comentropy related to original probabilities *p* in all bins.

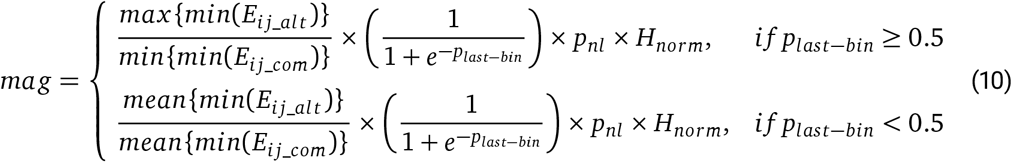

where

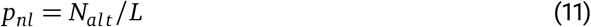

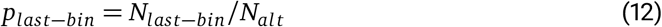

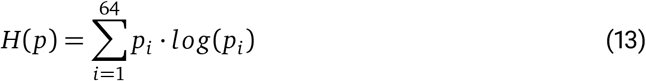

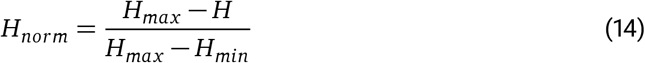

The final energy function for common distributions is: *E*_*ij_com*_ *final*_ = *mag E*_*i*j_*com*_, where *mag* is calculated based on the minimum energies of alternative and common signals, adjusted by *p*_*nl*_, *p*_*last*−*bin*_, and *H*_*norm*_.

This approach allows us to systematically model alternative protein conformations while accounting for the complex interplay between different structural signals. The opening or widening tendency between domains or helices is normally harder to achieve. By differentiating between alternative and common signals and applying appropriate scaling, we aim to achieve more accurate predictions of the alternative conformation.

#### 4.5.2 Assessment of modeled structures

We evaluated the modeled structures using Rosetta’s energy score function, which comprises two components: *E*_*Rosetta*_ and *E*_*Restraint*_. *E*_*Rosetta*_ is the sum of all energy terms in the *ref2015* scoring function, ^50^ commonly used to assess the plausibility of Rosetta-built structures. Negative *E*_*Rosetta*_ values indicate plausible structures. *E*_*Restraint*_ represents the energy resulting from distance restraints between all residue pairs, specifically using the *AtomPair* restraint for distance-based potentials (*E*_*ij_alt*_, *E*_*ij_com*_ *final*_). We only considered models with negative *E*_*Restraint*_ for further analysis, as positive values indicate failure to meet predicted distance restraints.

For each protein, we normalized the energy scores across 150 models sampled in one prediction using Eq. (15). *i* represents each modeled structure, and *E*_*Rosetta_max*_ and *E*_*Rosetta_min*_ are the maximum and minimum energy scores, respectively.

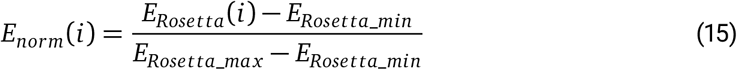

Model Selection Criteria:

From the perspective of prediction, we developed a set of criteria to select the best-scored structures based on energy scores and TM-score differences (Δ*T M*) relative to the initial structure.

(1) Eliminate the 10% of models with the worst *E*_*Restraint*_ scores.
(2) Retain models with *E*_*Rosetta*_ *<* 0.
(3) Rank models by *E*_*norm*_, focusing on those with *E*_*norm*_ *<* 0.8 * *max* (*E*_*norm*_).
(4) Calculate Δ*T M* (Δ*T M* = 1 − *T M*_*initial*−*relative*_) and select the top 10 models with the largest Δ*T M*.
(5) Choose the model with the lowest *E*_*norm*_ among these 10 as the final best-scored structure.

It is worth noting that these criteria were developed using proteins ALBP, DAHPS, and ZnT8 as model systems, with the remaining proteins used for split testing to avoid parameter overfitting.

We used RMSD and TM-score to assess the structural similarity between predicted models and the alternative structures solved experimentally. RMSD was calculated using the equation 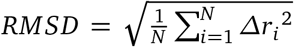 after aligning the *C*_α_ atoms of the two structures, where Δ*r*_*i*_ is the distance between each pair of corresponding atoms. TM-score was calculated using TM-Tools developed by Zhang Lab ^51^ to describe the topological similarity of conformations. Visualization of structures was performed using PyMOL (https://pymol.org/).

## Supporting information

codes and demos

## 5 Data availability

All the structural data sets and input files used in this study are provided in the supplementary files.

## 6 Code availability

All the source code and scripts for modeling and analyses are provided in the supplementary files.

## 7 Acknowledgement

This work was supported by the National Key R&D Program of China (2021YFE0108100) and the Science Fund for Creative Research Groups of the National Natural Science Foundation of China (T2321001).

## 8 Author contributions

C.S. designed and supervised the project. J.L. conducted the research, including the DM analysis and structural modeling. Z.Z. contributed to data analysis. All the authors participated in the writing of the manuscript.

## 9 Competing interests

The authors declare no competing interests.

## Supplementary Information

### Supplementary Tables

**Table S1.** Details of the 20 protein systems analyzed in this study. PDB IDs for the experimental structures of conformation 1 (C1) and conformation 2 (C2) are provided. The penultimate column lists the sequence lengths of the aligned structures for the two states. The final column summarizes the types of conformational changes.

| Protein system | Name | C1_ID | C2_ID | Aligned sequence length | Type of conformational change |
| --- | --- | --- | --- | --- | --- |
| **T4L** | T4 Lysozyme | 172L_A | 2LZM_A | 161 | hinge-bending |
| **AK** | Adenylate kinase | 4AKE_B | 2ECK_B | 214 | combination of changes in multi-domains |
| **LBP** | Leucine binding protein | 1USG_A | 1USI_A | 345 | hinge-bending |
| **ALBP** | D-allose binding protein | 1GUD_A | 1RPJ_A | 288 | hinge-bending |
| **RBP** | Ribose binding protein | 1URP_A | 2DRI_A | 271 | hinge-bending |
| **GroEL** | 60 kDa chaperonin | 1AON_A | 4HEL_A | 524 | combination of changes in multi-domains |
| **GlnBP** | Glutamine binding protein | 1GGG_A | 1WDN_A | 226 | hinge-bending |
| **STP10** | Sugar transport protein 10 | 7AAQ_A | 7AAR_A | 480 | transmembrane helices & loops |
| **DAHPS** | DAHPS synthase | 1VR6_A | 1RZM_A | 338 | domain rotation |
| **S100B** | S100 calcium-binding protein | 1SYM-2_B | 1XYD-15_A | 83 | single domain helices & loops |
| **CNBD** | Nucleotide-activated K<sup>+</sup> channel binding protein | 2KXL-8_A | 2K0G-9_A | 142 | single domain helices & loops |
| **HSPD1** | Human mitochondrial chaperonin | 7AZP_A | 4PJ1_A | 524 | combination of changes in multi-domains |
| **NTnC** | Regulatory domain of Troponin C | 1SKT-10_A | 1TNQ-33_A | 90 | single domain helices & loops |
| **ZnT8** | Zinc transporter 8 | 6XPF_A | 6XPD_A | 194 | transmembrane helices & loops |
| **NTRC** | Receiver domain of transcriptional enhancer binding protein | 1NTR-7_A | 1KRX-4_A | 124 | single domain helices & loops |
| **Mhp1** | Hydantoin transport protein | 2JLN_A | 2X79_A | 465 | transmembrane helices & loops |
| **ASCT2** | Neutral amino acid transporter | 6GCT_A | 7BCQ_A | 427 | transmembrane helices & loops |
| **AP4A** | Ap4A hydrolase | 1F3Y-17_A | 1JKN-11_A | 165 | single domain helices & loops |
| **S100A1** | S100 metal binding protein | 1K2H-2_A | 1ZFS-13_B | 93 | single domain helices & loops |
| **48G7g** | Immunoglobulin 48G7 germline Fab | 2RCS_H | 1AJ7_H | 217 | domain rotation |

**Table S2.** Evaluation of control experiment 1 (CTRL1) using the entire distance map predicted by the default AF2 for modeling the 20 selected proteins. The superscripts, C1 and C2, indicate the conformations relative to which the TM-scores were calculated. Successful predictions are highlighted in bold.

| Protein | TM<sub>2states</sub> | TM-scores (C1 to C2) |  |  |  | TM-scores (C2 to C1) |  |  |  |
| --- | --- | --- | --- | --- | --- | --- | --- | --- | --- |
|  |  | Best<sub>C1</sub><br>scored | Best<sub>C2</sub><br>scored | Best<sub>C1</sub><br>sampled | Best<sub>C2</sub><br>sampled | Best<sub>C1</sub><br>scored | Best<sub>C2</sub><br>scored | Best<sub>C1</sub><br>sampled | Best<sub>C2</sub><br>sampled |
| AK | **0.69** | 0.74 | **0.90** | 0.73 | **0.91** | 0.74 | 0.90 | 0.74 | 0.90 |
| LBP | **0.64** | 0.86 | 0.75 | 0.83 | 0.77 | 0.66 | 0.99 | 0.67 | 0.99 |
| ALBP | **0.76** | 0.78 | **0.95** | 0.79 | **0.95** | 0.78 | 0.95 | 0.79 | 0.95 |
| RBP | **0.76** | 0.78 | **0.99** | 0.78 | **0.99** | 0.78 | 0.99 | 0.79 | 0.99 |
| GroEL | **0.64** | 0.64 | **0.78** | 0.64 | **0.82** | 0.66 | 0.75 | 0.69 | 0.77 |
| GlnBP | **0.67** | 0.84 | 0.70 | 0.86 | 0.71 | 0.66 | 0.98 | 0.67 | 0.98 |
| SPT10 | **0.82** | 0.84 | **0.93** | 0.88 | **0.93** | 0.84 | 0.95 | 0.85 | 0.95 |
| DAHPS | **0.81** | 0.84 | 0.81 | 0.84 | 0.81 | **0.83** | 0.81 | **0.84** | 0.81 |
| S100B | **0.55** | 0.60 | **0.82** | 0.60 | **0.82** | 0.60 | 0.84 | 0.61 | 0.84 |
| CNBD | **0.74** | 0.71 | **0.86** | 0.71 | **0.87** | 0.71 | 0.98 | 0.71 | 0.85 |
| NTRC | **0.66** | 0.68 | **0.75** | 0.68 | **0.76** | 0.68 | 0.75 | 0.69 | 0.75 |
| AP4A | **0.80** | 0.84 | **0.86** | 0.83 | **0.87** | 0.83 | 0.87 | 0.83 | 0.87 |
| 48G7g | **0.57** | 0.57 | **0.81** | 0.57 | **0.82** | 0.60 | 0.79 | 0.62 | 0.80 |
| S100A1 | **0.50** | 0.51 | **0.72** | 0.52 | **0.73** | 0.51 | 0.73 | 0.53 | 0.74 |
| T4L | **0.76** | 0.97 | 0.80 | 0.97 | 0.81 | **0.97** | 0.80 | **0.97** | 0.80 |
| NTnC | **0.55** | 0.63 | **0.74** | 0.62 | **0.74** | 0.63 | 0.72 | 0.64 | 0.73 |
| HSPD1 | **0.64** | 0.66 | **0.74** | 0.67 | **0.75** | 0.62 | 0.78 | 0.64 | 0.80 |
| ZnT8 | **0.70** | 0.83 | 0.82 | 0.83 | 0.82 | **0.83** | 0.82 | **0.83** | 0.81 |
| Mhp1 | **0.88** | 0.96 | 0.89 | 0.96 | 0.89 | **0.95** | 0.88 | **0.96** | 0.89 |
| ASCT2 | **0.65** | 0.63 | 0.53 | 0.97 | 0.65 | 0.56 | 0.55 | 0.67 | 0.82 |

**Table S3.** Evaluation of control experiment 2 (CTRL2) without using the scaling parameter ‘mag’ in the energy functions for modeling the 20 selected proteins. The superscripts, C1 and C2, indicate the conformations relative to which the TM-scores were calculated. Successful predictions are highlighted in bold.

| Protein | TM<sub>2states</sub> | TM-scores (C1 to C2) |  |  |  | TM-scores (C2 to C1) |  |  |  |
| --- | --- | --- | --- | --- | --- | --- | --- | --- | --- |
|  |  | Best<sub>C1</sub><br>scored | Best<sub>C2</sub><br>scored | Best<sub>C1</sub><br>sampled | Best<sub>C2</sub><br>sampled | Best<sub>C1</sub><br>scored | Best<sub>C2</sub><br>scored | Best<sub>C1</sub><br>sampled | Best<sub>C2</sub><br>sampled |
| AK | **0.69** | 0.72 | **0.91** | 0.72 | **0.92** | 0.77 | 0.87 | 0.79 | 0.85 |
| LBP | **0.64** | 0.84 | 0.78 | 0.78 | **0.84** | 0.67 | 0.99 | 0.67 | 0.99 |
| ALBP | **0.76** | 0.78 | **0.95** | 0.79 | **0.95** | 0.79 | 0.95 | 0.79 | 0.95 |
| RBP | **0.76** | 0.78 | **0.99** | 0.78 | **0.99** | 0.78 | 0.99 | 0.78 | 0.99 |
| GroEL | **0.64** | 0.78 | 0.63 | 0.82 | 0.64 | 0.78 | 0.69 | 0.78 | 0.70 |
| GlnBP | **0.67** | 0.85 | 0.72 | 0.83 | 0.74 | 0.66 | 0.98 | 0.67 | 0.98 |
| SPT10 | **0.82** | 0.90 | **0.93** | 0.90 | **0.93** | 0.92 | 0.91 | 0.92 | 0.91 |
| DAHPS | **0.81** | 0.83 | 0.81 | 0.84 | 0.81 | **0.83** | 0.81 | **0.85** | 0.81 |
| S100B | **0.55** | 0.60 | **0.83** | 0.60 | **0.83** | 0.60 | 0.83 | 0.61 | 0.84 |
| CNBD | **0.74** | 0.71 | **0.87** | 0.71 | **0.87** | 0.71 | 0.84 | 0.71 | 0.85 |
| NTRC | **0.66** | 0.68 | **0.75** | 0.68 | **0.76** | 0.68 | 0.75 | 0.69 | 0.75 |
| AP4A | **0.80** | 0.83 | **0.88** | 0.83 | **0.88** | 0.83 | 0.87 | 0.83 | 0.87 |
| 48G7g | **0.57** | 0.57 | **0.81** | 0.57 | **0.84** | 0.59 | 0.79 | 0.61 | 0.80 |
| S100A1 | **0.50** | 0.51 | **0.73** | 0.51 | **0.74** | 0.51 | 0.75 | 0.52 | 0.74 |
| T4L | **0.76** | 0.96 | 0.80 | 0.97 | 0.81 | **0.96** | 0.80 | **0.97** | 0.80 |
| NTnC | **0.55** | 0.63 | **0.72** | 0.62 | **0.74** | 0.63 | 0.73 | 0.64 | 0.73 |
| HSPD1 | **0.64** | 0.66 | **0.71** | 0.66 | **0.75** | 0.64 | 0.78 | 0.64 | 0.78 |
| ZnT8 | **0.70** | 0.82 | **0.83** | 0.82 | **0.83** | **0.83** | 0.79 | **0.83** | 0.80 |
| Mhp1 | **0.88** | 0.96 | 0.90 | 0.95 | 0.90 | **0.95** | 0.88 | **0.96** | 0.88 |
| ASCT2 | **0.65** | 0.62 | 0.53 | 0.98 | 0.65 | 0.57 | 0.55 | 0.68 | 0.77 |

### Supplementary Figures

**Figure S1.**
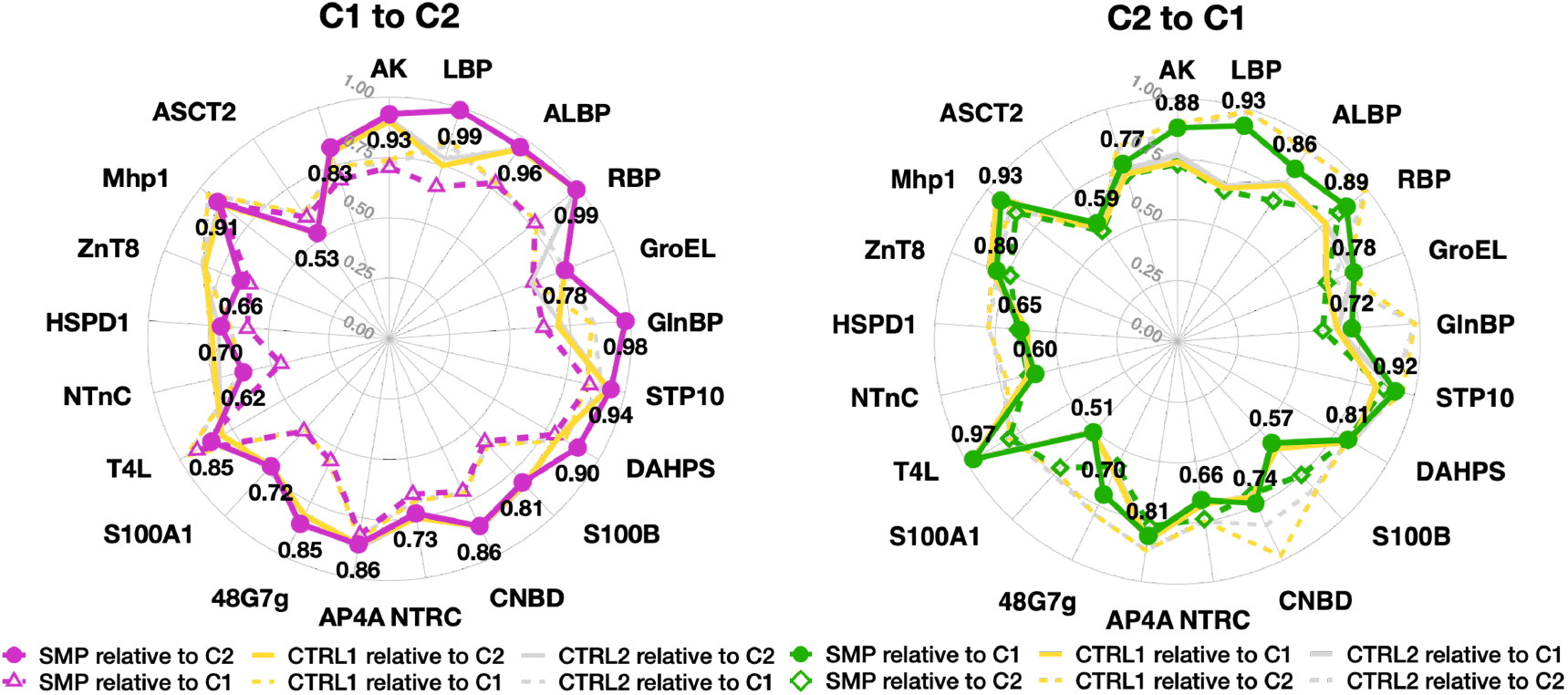
Radar maps comparing TM-scores of best-scored samples (SMP) with two control experiments. Control experiment 1 (CTRL1, yellow): sampling with the entire AF2-predicted distance map; control experiment 2 (CTRL2, grey): sampling without the scaling parameter ‘mag’ in energy functions. Solid lines represent TM-scores relative to the target state; dotted lines indicate TM-scores relative to the initial known state. Effective prediction rates for the second state are 85% (C1 to C2) and 70% (C2 to C1), where predicted models are more similar to the target state and further from the initial state. Control experiments show lower success rates: 65% and 15% for the two directions, respectively. The peripheral distribution of dotted lines indicates that structures often remain close to the initial state in the control sampling.

**Figure S2.**
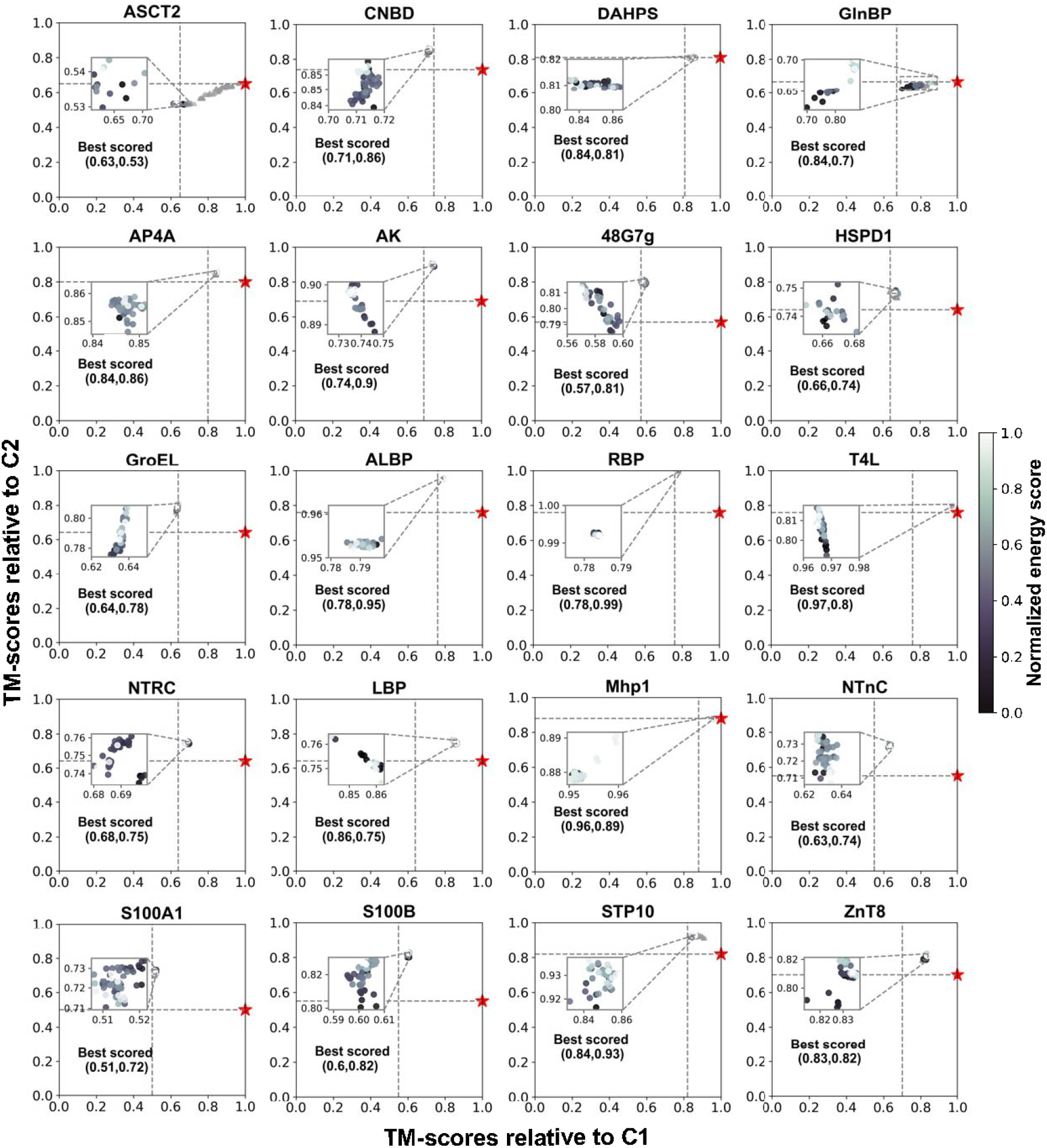
Rosetta Sampling Results for Control Experiment 1 (CTRL1). The figure displays the outcomes of Rosetta sampling using the full distance map predicted by default AlphaFold2 (AF2) from C1 to C2. The initial structure is marked with a red star, while 150 generated models are color-coded according to their normalized energy scores (darker colors represent lower energy). The dotted line illustrates the initial conformational similarity between states (TM_2*states*_). Grey triangles indicate structures that do not meet the designed restraints. The inset provides a zoomed-in view of clustered models, highlighting the best-scored model, which is the predicted structure with the lowest energy score. The TM-scores relative to C1 and C2 are shown in each panel.

**Figure S3.**
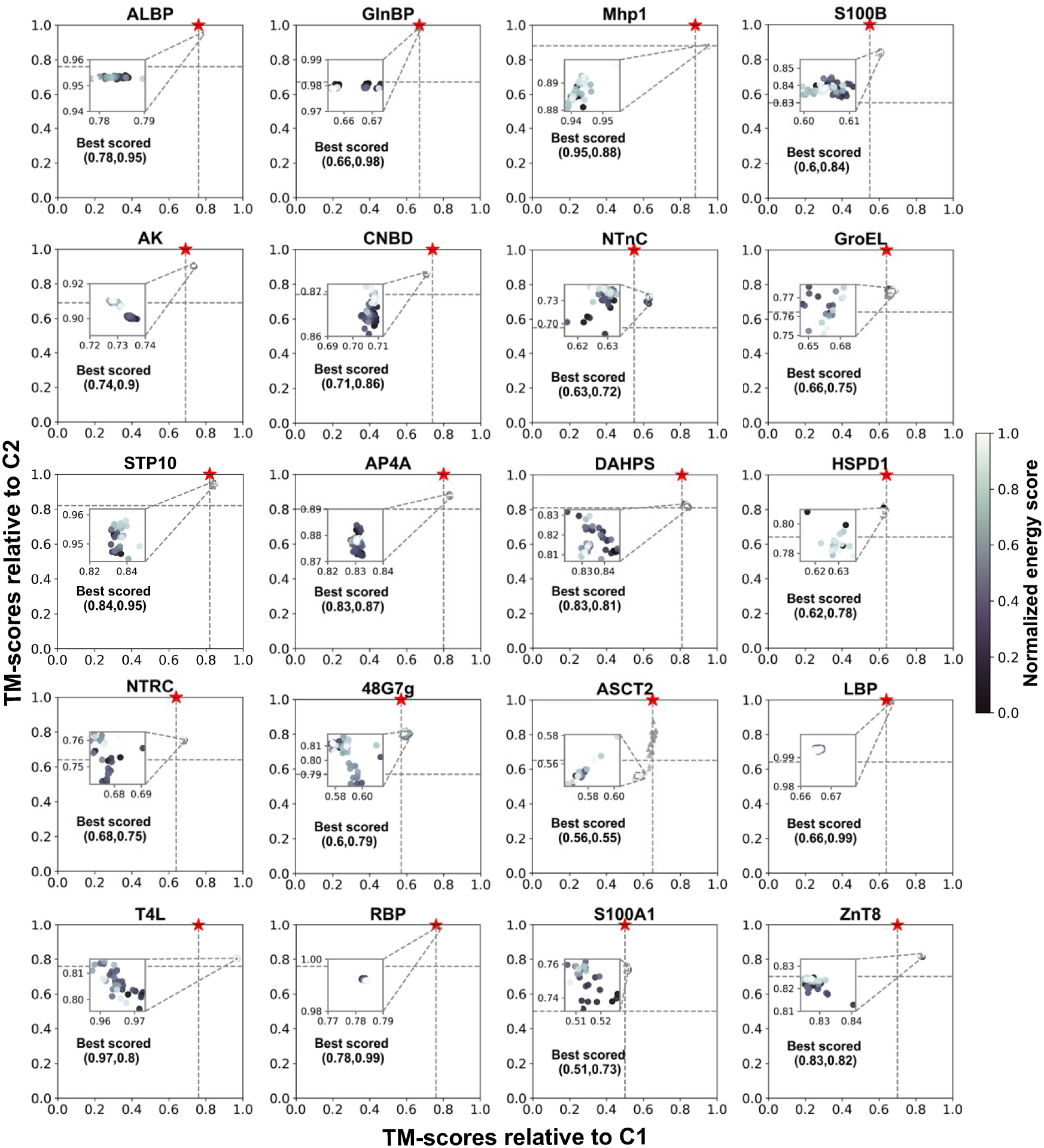
Rosetta Sampling Results for Control Experiment 1 (CTRL1). The figure displays the outcomes of Rosetta sampling using the full distance map predicted by default AlphaFold2 (AF2) from C2 to C1. The initial structure is marked with a red star, while 150 generated models are color-coded according to their normalized energy scores (darker colors represent lower energy). The dotted line illustrates the initial conformational similarity between states (TM_2*states*_). Grey triangles indicate structures that do not meet the designed restraints. The inset provides a zoomed-in view of clustered models, highlighting the best-scored model, which is the predicted structure with the lowest energy score. The TM-scores relative to C1 and C2 are shown in each panel.

**Figure S4.**
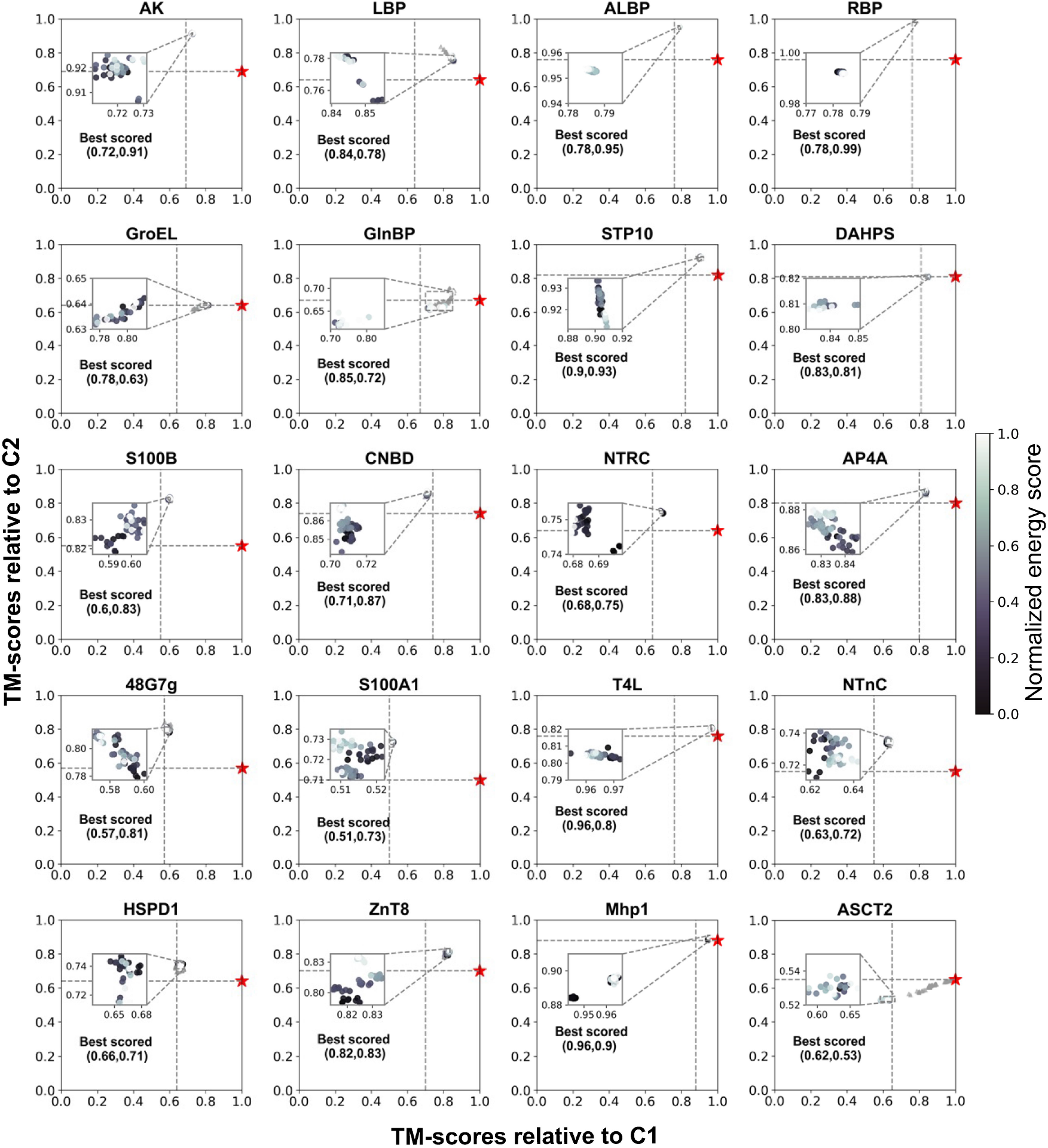
Rosetta Sampling Results for Control Experiment 2 (CTRL2). This figure presents the results of Rosetta sampling conducted without applying the scaling factor ‘mag’ in the energy functions during the transition from C1 to C2. The initial structure is denoted by a red star, and 150 generated models are displayed, with colors representing their normalized energy scores (darker colors indicate lower energy). The dotted line signifies the initial conformational similarity between states (TM_2*states*_). Grey triangles denote structures that failed to satisfy the designed restraints. The inset offers a zoomed-in view of the clustered models, with a focus on the best-scored model – the predicted structure with the lowest energy score. The TM-scores relative to C1 and C2 are shown in each panel.

**Figure S5.**
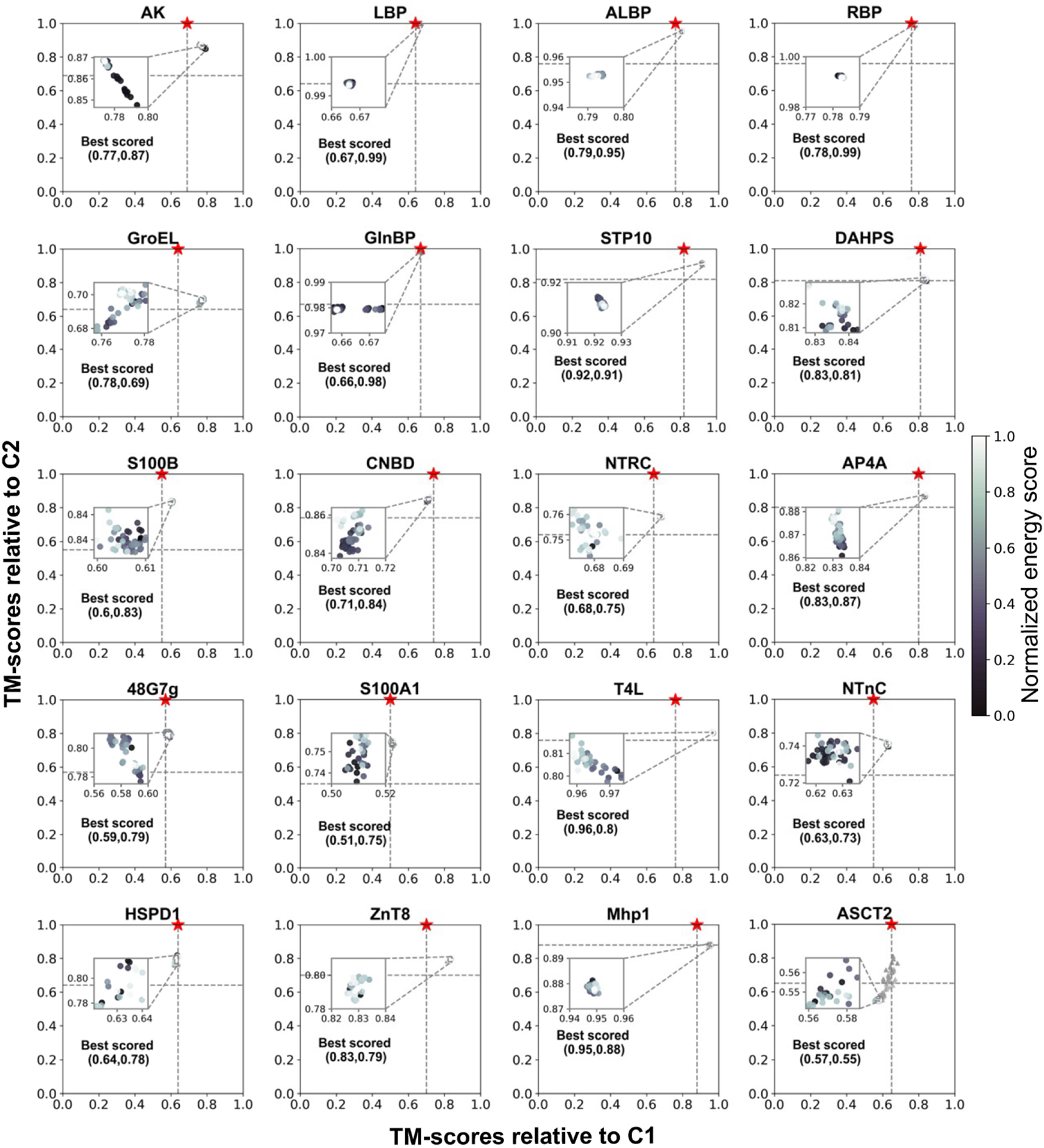
Rosetta Sampling Results for Control Experiment 2 (CTRL2). This figure presents the results of Rosetta sampling conducted without applying the scaling factor ‘mag’ in the energy functions during the transition from C2 to C1. The initial structure is denoted by a red star, and 150 generated models are displayed, with colors representing their normalized energy scores (darker colors indicate lower energy). The dotted line signifies the initial conformational similarity between states (TM_2*states*_). Grey triangles denote structures that failed to satisfy the designed restraints. The inset offers a zoomed-in view of the clustered models, with a focus on the best-scored model – the predicted structure with the lowest energy score. The TM-scores relative to C1 and C2 are shown in each panel.

**Figure S6.**
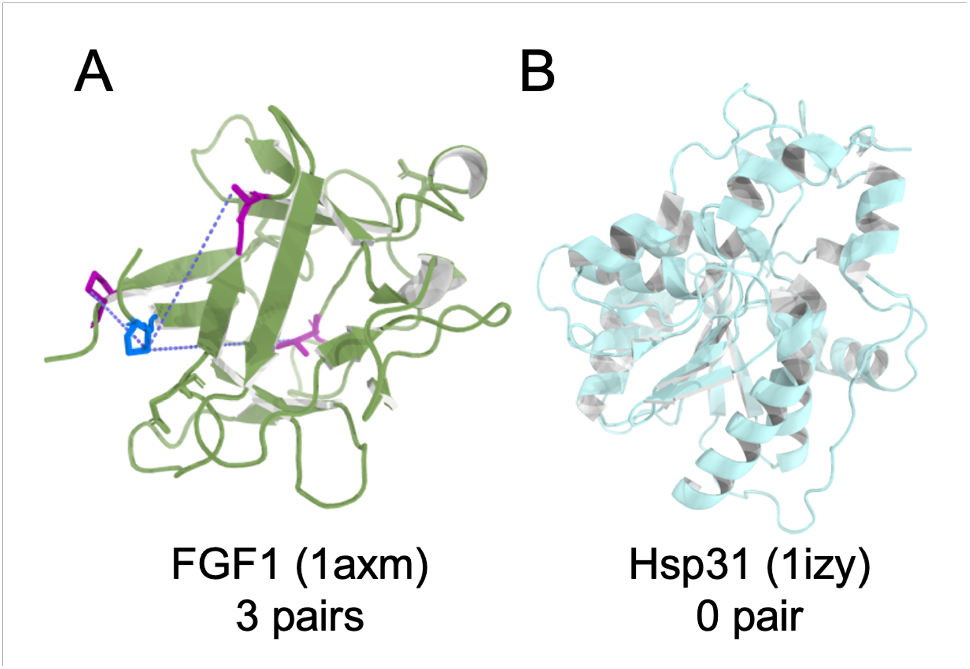
Structures and predicted signals for two extremostable proteins. (A) Fibroblast growth factor 1 (FGF1, PDB ID: 1axm): three signals detected, connected with loops at sequence end. (B) Heat shock protein 31 (Hsp31, PDB ID: 1izy): no alternative signals predicted.

## Notes

### Competing Interest Statement

The authors have declared no competing interest.

### Summary of Updates

change the article category into New Results

