## Supplementary material for "Predicting the alternative conformation of a known protein structure based on the distance map of AlphaFold2": codes and demos: msa.sh

```
#!/bin/bash -e
MMSEQS="$1"
QUERY="$2"
BASE="$4"
DB1="$5"
DB2="$6"
DB3="$7"
USE_ENV="$8"
USE_TEMPLATES="$9"
FILTER="${10}"
TAXONOMY="${11}"
M8OUT="${12}"
EXPAND_EVAL=inf
ALIGN_EVAL=10
DIFF=3000
QSC=-20.0
MAX_ACCEPT=1000000
if [ "${FILTER}" = "1" ]; then
# 0.1 was not used in benchmarks due to POSIX shell bug in line above
#  EXPAND_EVAL=0.1
  ALIGN_EVAL=10
  QSC=0.8
  MAX_ACCEPT=100000
fi
export MMSEQS_CALL_DEPTH=1
SEARCH_PARAM="--num-iterations 3 --db-load-mode 2 -a --k-score 'seq:96,prof:80' -e 0.1 --max-seqs 10000"
FILTER_PARAM="--filter-min-enable 1000 --diff ${DIFF} --qid 0.0,0.2,0.4,0.6,0.8,1.0 --qsc 0 --max-seq-id 0.95"
EXPAND_PARAM="--expansion-mode 0 -e ${EXPAND_EVAL} --expand-filter-clusters ${FILTER} --max-seq-id 0.95"
mkdir -p "${BASE}"
"${MMSEQS}" createdb "${QUERY}" "${BASE}/qdb"
"${MMSEQS}" search "${BASE}/qdb" "${DB1}" "${BASE}/res" "${BASE}/tmp1" $SEARCH_PARAM
"${MMSEQS}" mvdb "${BASE}/tmp1/latest/profile_1" "${BASE}/prof_res"
"${MMSEQS}" lndb "${BASE}/qdb_h" "${BASE}/prof_res_h"

(

"${MMSEQS}" expandaln "${BASE}/qdb" "${DB1}.idx" "${BASE}/res" "${DB1}.idx" "${BASE}/res_exp" --db-load-mode 2 ${EXPAND_PARAM}
"${MMSEQS}" align "${BASE}/prof_res" "${DB1}.idx" "${BASE}/res_exp" "${BASE}/res_exp_realign" --db-load-mode 2 -e ${ALIGN_EVAL} --max-accept ${MAX_ACCEPT} --alt-ali 10 -a
"${MMSEQS}" filterresult "${BASE}/qdb" "${DB1}.idx" "${BASE}/res_exp_realign" "${BASE}/res_exp_realign_filter" --db-load-mode 2 --qid 0 --qsc $QSC --diff 0 --max-seq-id 1.0 --filter-min-enable 100
if [ "${M8OUT}" = "1" ]; then
  "${MMSEQS}" filterresult "${BASE}/qdb" "${DB1}.idx" "${BASE}/res_exp_realign_filter" "${BASE}/res_exp_realign_filter_filter" --db-load-mode 2 ${FILTER_PARAM}
  "${MMSEQS}" convertalis "${BASE}/qdb" "${DB1}.idx" "${BASE}/res_exp_realign_filter_filter" "${BASE}/uniref.m8" --db-load-mode 2 --format-output query,target,fident,alnlen,mismatch,gapopen,qstart,qend,tstart,tend,evalue,bits,tseq
  "${MMSEQS}" rmdb "${BASE}/res_exp_realign_filter_filter"
else
  "${MMSEQS}" result2msa "${BASE}/qdb" "${DB1}.idx" "${BASE}/res_exp_realign_filter" "${BASE}/uniref.a3m" --msa-format-mode 6 --db-load-mode 2 --filter-msa ${FILTER} ${FILTER_PARAM}
fi
"${MMSEQS}" rmdb "${BASE}/res_exp_realign"
"${MMSEQS}" rmdb "${BASE}/res_exp"
"${MMSEQS}" rmdb "${BASE}/res"
if [ "${TAXONOMY}" = "1" ] && [ -e "${DB1}_taxonomy" ]; then
  "${MMSEQS}" convertalis "${BASE}/qdb" "${DB1}.idx" "${BASE}/res_exp_realign_filter" "${BASE}/res_exp_realign_tax" --db-output 1 --format-output "taxid,target,taxlineage" --db-load-mode 2
  awk 'BEGIN { printf("%c%c%c%c",8,0,0,0); exit; }' > "${BASE}/res_exp_realign_tax.dbtype"
  MMSEQS_FORCE_MERGE=1 "${MMSEQS}" filtertaxdb "${DB1}" "${BASE}/res_exp_realign_tax" "${BASE}/res_exp_realign_tax_filt" --taxon-list '!12908&&!28384'
  tr -d '\000' < "${BASE}/res_exp_realign_tax_filt" | sort -u > "${BASE}/uniref_tax.tsv"
fi
"${MMSEQS}" rmdb "${BASE}/res_exp_realign_filter"

)&
(

if [ "${USE_TEMPLATES}" = "1" ]; then
  "${MMSEQS}" search "${BASE}/prof_res" "${DB2}" "${BASE}/res_pdb" "${BASE}/tmp2" --db-load-mode 2 -s 7.5 -a -e 0.1
  "${MMSEQS}" convertalis "${BASE}/prof_res" "${DB2}.idx" "${BASE}/res_pdb" "${BASE}/pdb70.m8" --format-output query,target,fident,alnlen,mismatch,gapopen,qstart,qend,tstart,tend,evalue,bits,cigar --db-load-mode 2
  "${MMSEQS}" rmdb "${BASE}/res_pdb"
fi

)&
(

if [ "${USE_ENV}" = "1" ]; then
  "${MMSEQS}" search "${BASE}/prof_res" "${DB3}" "${BASE}/res_env" "${BASE}/tmp3" $SEARCH_PARAM
  "${MMSEQS}" expandaln "${BASE}/prof_res" "${DB3}.idx" "${BASE}/res_env" "${DB3}.idx" "${BASE}/res_env_exp" -e ${EXPAND_EVAL} --expansion-mode 0 --db-load-mode 2
  "${MMSEQS}" align "${BASE}/tmp3/latest/profile_1" "${DB3}.idx" "${BASE}/res_env_exp" "${BASE}/res_env_exp_realign" --db-load-mode 2 -e ${ALIGN_EVAL} --max-accept ${MAX_ACCEPT} --alt-ali 10 -a
  "${MMSEQS}" filterresult "${BASE}/qdb" "${DB3}.idx" "${BASE}/res_env_exp_realign" "${BASE}/res_env_exp_realign_filter" --db-load-mode 2 --qid 0 --qsc $QSC --diff 0 --max-seq-id 1.0 --filter-min-enable 100
  if [ "${M8OUT}" = "1" ]; then
    "${MMSEQS}" filterresult "${BASE}/qdb" "${DB3}.idx" "${BASE}/res_env_exp_realign_filter" "${BASE}/res_env_exp_realign_filter_filter" --db-load-mode 2 ${FILTER_PARAM}
    "${MMSEQS}" convertalis "${BASE}/qdb" "${DB3}.idx" "${BASE}/res_env_exp_realign_filter_filter" "${BASE}/bfd.mgnify30.metaeuk30.smag30.m8" --db-load-mode 2 --format-output query,target,fident,alnlen,mismatch,gapopen,qstart,qend,tstart,tend,evalue,bits,tseq
    "${MMSEQS}" rmdb "${BASE}/res_env_exp_realign_filter_filter"
  else
	"${MMSEQS}" result2msa "${BASE}/qdb" "${DB3}.idx" "${BASE}/res_env_exp_realign_filter" "${BASE}/bfd.mgnify30.metaeuk30.smag30.a3m" --msa-format-mode 6 --db-load-mode 2 --filter-msa ${FILTER} ${FILTER_PARAM}
  fi
  "${MMSEQS}" rmdb "${BASE}/res_env_exp_realign_filter"
  "${MMSEQS}" rmdb "${BASE}/res_env_exp_realign"
  "${MMSEQS}" rmdb "${BASE}/res_env_exp"
  "${MMSEQS}" rmdb "${BASE}/res_env"
fi

)&
wait

"${MMSEQS}" rmdb "${BASE}/qdb"
"${MMSEQS}" rmdb "${BASE}/qdb_h"
"${MMSEQS}" rmdb "${BASE}/res"
rm -f -- "${BASE}/prof_res"*
rm -rf -- "${BASE}/tmp1" "${BASE}/tmp2" "${BASE}/tmp3"
```
