## Supplementary material for "Predicting the alternative conformation of a known protein structure based on the distance map of AlphaFold2": codes and demos: arguments_AF2.py

```
import argparse

def get_args(params):

    parser = argparse.ArgumentParser(formatter_class=argparse.ArgumentDefaultsHelpFormatter)
    parser.add_argument("FASTA", type=str, help="input sequence")
    #parser.add_argument("NPZ", type=str, help="npz file of saved distance...")
    parser.add_argument("OUT", type=str, help="output model (in PDB format)")
    parser.add_argument('-pd', type=float, dest='pcut', default=params['PCUT'], help='min probability of distance restraints')
    parser.add_argument('-m', type=int, dest='mode', default=2, choices=[0,1,2], help='0: sh+m+l, 1: (sh+m)+l, 2: (sh+m+l)')
    parser.add_argument('-w', type=str, dest='wdir', default=params['WDIR'], help='folder to store temp files')
    parser.add_argument('-n', type=int, dest='steps', default=1000, help='number of minimization steps')
    parser.add_argument('--orient', dest='use_orient', action='store_true', help='use orientations')
    parser.add_argument('--no-orient', dest='use_orient', action='store_false')
    parser.add_argument('--fastrelax', dest='fastrelax', action='store_true', help='perform FastRelax')
    parser.add_argument('--no-fastrelax', dest='fastrelax', action='store_false')
    #parser.set_defaults(use_orient=True)
    parser.set_defaults(use_orient=False)
    parser.set_defaults(fastrelax=True)

    args = parser.parse_args()

    params['PCUT'] = args.pcut
    params['USE_ORIENT'] = args.use_orient

    return args
```
