## Supplementary material for "Predicting the alternative conformation of a known protein structure based on the distance map of AlphaFold2": codes and demos: dij_magH_noroll_Utils_cycle_ros_dist_npy.py

```
import numpy as np
import random
from pyrosetta import *
import math
from scipy import stats
from scipy.special import expit
np.set_printoptions(threshold=200)

eps =1e-9

def gen_rst(npz, u, dist_flag,H, tmpdir, params,pose,p_last,p_nl):

    dist = npz
    ##Apo / Holo flag?!
    num = len(np.where(list(dist_flag))[0])  
    flag = np.where(list(dist_flag))
    print("diff DM points:",num,flag,len(dist[0]))
    #dist_index = np.zeros(dist.shape)
    
    #print("after flag",dist_holo[flag[0][2]][flag[1][2]])
 
    #dist = np.roll(dist,1,axis=-1)
    dist = dist.astype(np.float32)+eps
    #print("original dist",dist[14,189])
    # dictionary to store Rosetta restraints
    rst = {'dist' : [], 'omega' : [], 'theta' : [], 'phi' : [], 'rep' : []}

    ########################################################
    # assign parameters
    ########################################################
    #PCUT  = 0.95 #params['PCUT']
    PCUT  = 0.0005 #params['PCUT']
    PCUT1 = params['PCUT1']
    EBASE = params['EBASE']
    EREP  = params['EREP']
    DREP  = params['DREP']
    PREP  = params['PREP']
    SIGD  = params['SIGD']
    SIGM  = params['SIGM']
    MEFF  = params['MEFF']
    #DCUT  = params['DCUT']
    DCUT = 21.6875
    ALPHA = params['ALPHA']
    MEFF=0.0001
    EBASE=-0.5

    #DSTEP = params['DSTEP']
    DSTEP = 0.3125
    ASTEP = np.deg2rad(params['ASTEP'])

    seq = params['seq']

    ########################################################
    # repultion restraints
    ########################################################
    #cbs = ['CA' if a=='G' else 'CB' for a in params['seq']]
    '''
    prob = np.sum(dist[:,:,5:], axis=-1)
    i,j = np.where(prob<PREP)
    prob = prob[i,j]
    for a,b,p in zip(i,j,prob):
        if b>a:
            name=tmpdir.name+"/%d.%d_rep.txt"%(a+1,b+1)
            rst_line = 'AtomPair %s %d %s %d SCALARWEIGHTEDFUNC %.2f SUMFUNC 2 CONSTANTFUNC 0.5 SIGMOID %.3f %.3f\n'%('CB',a+1,'CB',b+1,-0.5,SIGD,SIGM)
            rst['rep'].append([a,b,p,rst_line])
    print("rep restraints:   %d"%(len(rst['rep'])))
    '''
    ########################################################
    # dist: 0..20A
    ########################################################
    nres = dist.shape[0]
    ###original p<PCUT abandoned
    #for c in range(0,nres):
        #for d in range(0,nres):
            #vec = np.where(dist[c,d]>(PCUT+eps))
            #print(vec[0],dist[c,d][vec[0]])
            #for e in range(0,64):
                #if e not in vec[0]:
                    #dist[c,d,e]=float(eps)
    bins = np.array([3.25+DSTEP*i for i in range(61)]) #DSTEP=0.3125
    prob = np.sum(dist[:,:,5:], axis=-1) #p for 3.875 A further
    bkgr = np.array((bins/DCUT)**ALPHA) #DCUT=21.6875, ALPHA = 1.57
    print('original distInput',dist[57,177])
    #attr = -np.log((dist[:,:,5:]+MEFF)/(dist[:,:,-1][:,:,None]*bkgr[None,None,:]))+EBASE #MEFF=0.0001,EBASE=-0.5
    #attr = -np.log((dist[:,:,3:]+MEFF)/(np.sum(dist[:,:,-3:-1][:,:,None],axis=3)*bkgr[None,None,:]))+EBASE #MEFF=0.0001,EBASE=-0.5
    attr = -np.log((dist[:,:,3:]+MEFF)/(dist[:,:,-2][:,:,None]*bkgr[None,None,:]))+EBASE #MEFF=0.0001,EBASE=-0.5
    print('attr?',attr[57,177])
    repul = np.maximum(attr[:,:,0],np.zeros((nres,nres)))[:,:,None]+np.array(EREP)[None,None,:]  #EREP= [10.0,3.0,0.5]
    dist = np.concatenate([repul,attr], axis=-1)
    print('distProcessed?',dist[57,177])
    bins = np.concatenate([DREP,bins])  #DREP = [0.0,2.0,3.5]---0,2,3
    print("new bins",bins)
    i,j = np.where(prob>PCUT1)
    prob = prob[i,j]
    #test_attr = dist[:,:,-1][:,:,None]*bkgr[None,None,:] #only last bin for all pairs*bkgr [l,l,1]*[61]=[l,l,61]
    #print("bkgr:",bkgr,"dist-1*bkgr", test_attr.shape)
    inew = []
    jnew = []
    #print("i,j",i,j,len(i))
    #for n in range(0,len(i)):
        #x = np.where(dist[i[n],j[n]]<0)
        #if len(x[0])<=1 and int(x[0]) >=63:
            #print('d_distribution of last bin',i[n]+1,j[n]+1,len(x[0]),int(x[0]))
            #inew.append(i[n])
            #jnew.append(j[n])
            ##continue
        #else:
            #print('last bin',i[n]+1,j[n]+1,x[0])
            #inew.append(i[n])
            #jnew.append(j[n])
    #print("new i j:",len(inew),bins,len(bins))
    #prob = prob[inew,jnew]
    nbins = 64
    step = 0.3125
    #u = 0.1
    
    e_single = []
    min_edge = []
    max_edge = []
    min_edge_multi = []
    for a,b,p in zip(i,j,prob):
        if b>a+6:
            name=tmpdir.name+"/%d.%d.txt"%(a+1,b+1)
            tag = 0
            d_int = []
            resid_a=pose.residue(a+1)
            a_CB=resid_a.xyz("CB")
            resid_b=pose.residue(b+1)
            b_CB=resid_b.xyz("CB")
            a_b_vec= a_CB-b_CB
            dij=float('%.1f' %a_b_vec.norm())
            for n in range(0,num):
                if a==flag[0][n] and b==flag[1][n]:
                    #dist[a,b] = (1/(mag*H[a,b]))*dist[a,b]
                    tag = 1
                    for e in dist[a,b]:
                        if e>0:
                            d_int.append(int(e))
                    tag = 1
                    platform = stats.mode(d_int)[0][0]
                    if a==57 and b==177:
                        print("original",a+1,b+1,platform,'\n',dist[a,b])
                        print("Distance in ab:",dij)
                    for bin in range(0,64):
                        if dist[a,b,bin]>=0:
                            dist[a,b,bin] = math.tanh(dist[a,b,bin])
                    min_index =np.argmin(dist[a,b])
                    if dij<21.6875 and min_index>dij:
                        nleft = int((dij-2)/0.3125)
                        nright = np.where(dist[a,b]<0)[0][0]
                        print(nleft,nright,dist[a,b,nleft],dist[a,b,nright])
                        for bin in range(nleft,nright):
                            dist[a,b,bin]=dist[a,b,bin]-(1/(nright-nleft))*(bin-nleft)
                    elif dij<21.6875 and min_index<dij:
                        nright = int((dij-2)/0.3125)
                        nleft = np.where(dist[a,b]<0)[0][0]
                        print(nleft,nright,dist[a,b,nleft],dist[a,b,nright])
                        for bin in range(nleft,nright):
                            dist[a,b,bin]=dist[a,b,bin]-(1/(nright-nleft))*(bin-nleft)                       
                    #print("multi peaks_dist[a,b]",a+1,b+1,'\n',dist[a,b])
                    min_edge_multi.append(np.min(dist[a,b]))
                    #dist[a,b][np.where(dist[a,b]<0)[0]] = 20*dist[a,b][np.where(dist[a,b]<0)[0]] 
                    #if a==147:
                    #print("multi peaks_dist[a,b]",a+1,b+1,'\n',dist[a,b])
                    break
            if tag != 1:
                min_edge.append(np.min(dist[a,b]))
                d_int = []
                for ebasic in dist[a,b]:
                    if ebasic>0:
                        d_int.append(int(ebasic))
                platform = stats.mode(d_int)[0][0]
                #if a==147 and b==205:
                #print("single origin!!",a+1,b+1, platform,'\n',dist[a,b])
                for bin in range(0,64):
                    if dist[a,b,bin]>platform-1:
                    #if dist[a,b,bin]>=0:
                        dist[a,b,bin] = platform-1+math.tanh(dist[a,b,bin]-platform+1)
                e_single.append(np.mean(dist[a,b,2:]))
                max_edge.append(np.max(dist[a,b]))
                #print("single platform!!",a+1,b+1,'\n',dist[a,b])
                #print("e_single of 61bins",e_single[-1],a+1,b+1)

    min_multi = np.percentile(min_edge_multi,25)-1.5*(np.percentile(min_edge_multi,75)-np.percentile(min_edge_multi,25))
    #if max_multi>0:
        #max_multi = np.percentile(min_edge_multi,75)
    n_mini = 0
    for mini in min_edge_multi:
        if mini<min_multi:
            #print(mini)
            n_mini+=1
    #print("expabs_multi",min_edge_multi)
    #p_last = 0.66
    #p_nl = 0.82
    print("min_edge of original pairs:",min(min_edge),max(min_edge),"average",np.mean(min_edge),"percentile25,75",np.percentile(min_edge,25,interpolation='midpoint'),np.percentile(min_edge,75,interpolation='midpoint'))
    print("max_edge of original pairs:",min(max_edge),max(max_edge),"average",np.mean(max_edge),"percentile",np.percentile(max_edge,25,interpolation='midpoint'))
    print("everage of 61bins of single peak pairs:25/50/75percentile",np.percentile(e_single,25),np.percentile(e_single,50),np.percentile(e_single,75))
    print("min_edge_multi",min(min_edge_multi),max(min_edge_multi),"average",np.mean(min_edge_multi),"percentile25,75",np.percentile(min_edge_multi,25),np.percentile(min_edge_multi,75))
    print("1-sigmoid p",1-expit(p_last),"meanxp_nlx1-expit",p_nl*(1-expit(p_last))*(np.mean(min_edge_multi)/np.mean(min_edge)),'MaxMinxp_nlx1-expit',p_nl*(1-expit(p_last))*max(min_edge_multi)/min(min_edge))
    if p_last>0.5:
        #mag = p_nl*(1-expit(p_last))*(np.mean(min_edge_multi)/np.mean(min_edge))
        #mag =(1-expit(p_last))*(np.mean(min_edge_multi)/np.mean(min_edge))
        mag = p_nl*(1-expit(p_last))*max(min_edge_multi)/min(min_edge)
        #mag = (1-expit(p_last))*max(min_edge_multi)/min(min_edge)
        #mag = 0.001
    else:
        mag = p_nl*(1-expit(p_last))*(np.mean(min_edge_multi)/np.mean(min_edge))
        #mag = (1-expit(p_last))*(np.mean(min_edge_multi)/np.mean(min_edge))
        #mag = p_nl*(1-expit(p_last))*max(min_edge_multi)/min(min_edge)
        #mag = 1-expit(p_last)
        #mag = (1-expit(p_last))*max(min_edge_multi)/min(min_edge)
        #mag = 0.001
    print("mag in use:",mag,"max_edge/min_edge",max(min_edge_multi)/min(min_edge),"Mean",np.mean(min_edge_multi)/np.mean(min_edge),"min_multi",min_multi,n_mini)
    #for x in range(0,len(dist)):
        #for y in range(0,len(dist[0])):
            #if (x not in flag[0]) or (y not in flag[1]):
                #dist[x,y]=mag*dist[x,y]
                #print("rest pairs like",np.min(dist[x,y]))
    for a,b,p in zip(i,j,prob):
        if b>a+6:
            name=tmpdir.name+"/%d.%d.txt"%(a+1,b+1)
            dist[a,b]=(H[a,b]+1e-5)*mag*dist[a,b]
            #print(u*dist[a,b],a+1,b+1)
            for n in range(0,num):
                if a==flag[0][n] and b==flag[1][n]:
                    dist[a,b] = (1/(mag*(H[a,b]+1e-5)))*dist[a,b]
                    #print(dist[a,b],a+1,b+1)
                    break

            with open(name, "w") as f:
                f.write('x_axis'+'\t%.4f'*nbins%tuple(bins)+'\n')
                f.write('y_axis'+'\t%.4f'*nbins%tuple(u*dist[a,b])+'\n')
                f.close()
            rst_line = 'AtomPair %s %d %s %d SPLINE TAG %s 1.0 %.3f %.5f'%('CB',a+1,'CB',b+1,name,1.0,step)
            rst['dist'].append([a,b,p,rst_line])
            
    print("dist restraints:  %d"%(len(rst['dist'])))

    ########################################################
    # omega: -pi..pi
    ########################################################
    '''
    nbins = omega.shape[2]-1+4
    bins = np.linspace(-np.pi-1.5*ASTEP, np.pi+1.5*ASTEP, nbins)
    prob = np.sum(omega[:,:,1:], axis=-1)
    i,j = np.where(prob>PCUT)
    prob = prob[i,j]
    omega = -np.log((omega+MEFF)/(omega[:,:,-1]+MEFF)[:,:,None])
    omega = np.concatenate([omega[:,:,-2:],omega[:,:,1:],omega[:,:,1:3]],axis=-1)
    for a,b,p in zip(i,j,prob):
        if b>a:
            name=tmpdir.name+"/%d.%d_omega.txt"%(a+1,b+1)
            with open(name, "w") as f:
                f.write('x_axis'+'\t%.5f'*nbins%tuple(bins)+'\n')
                f.write('y_axis'+'\t%.5f'*nbins%tuple(omega[a,b])+'\n')
                f.close()
            rst_line = 'Dihedral CA %d CB %d CB %d CA %d SPLINE TAG %s 1.0 %.3f %.5f'%(a+1,a+1,b+1,b+1,name,1.0,ASTEP)
            rst['omega'].append([a,b,p,rst_line])
    print("omega restraints: %d"%(len(rst['omega'])))


    ########################################################
    # theta: -pi..pi
    ########################################################
    prob = np.sum(theta[:,:,1:], axis=-1)
    i,j = np.where(prob>PCUT)
    prob = prob[i,j]
    theta = -np.log((theta+MEFF)/(theta[:,:,-1]+MEFF)[:,:,None])
    theta = np.concatenate([theta[:,:,-2:],theta[:,:,1:],theta[:,:,1:3]],axis=-1)
    for a,b,p in zip(i,j,prob):
        if b!=a:
            name=tmpdir.name+"/%d.%d_theta.txt"%(a+1,b+1)
            with open(name, "w") as f:
                f.write('x_axis'+'\t%.3f'*nbins%tuple(bins)+'\n')
                f.write('y_axis'+'\t%.3f'*nbins%tuple(theta[a,b])+'\n')
                f.close()
            rst_line = 'Dihedral N %d CA %d CB %d CB %d SPLINE TAG %s 1.0 %.3f %.5f'%(a+1,a+1,a+1,b+1,name,1.0,ASTEP)
            rst['theta'].append([a,b,p,rst_line])
            #if a==0 and b==9:
            #    with open(name,'r') as f:
            #        print(f.read())
    print("theta restraints: %d"%(len(rst['theta'])))


    ########################################################
    # phi: 0..pi
    ########################################################
    nbins = phi.shape[2]-1+4
    bins = np.linspace(-1.5*ASTEP, np.pi+1.5*ASTEP, nbins)
    prob = np.sum(phi[:,:,1:], axis=-1)
    i,j = np.where(prob>PCUT)
    prob = prob[i,j]
    phi = -np.log((phi+MEFF)/(phi[:,:,-1]+MEFF)[:,:,None])
    phi = np.concatenate([np.flip(phi[:,:,1:3],axis=-1),phi[:,:,1:],np.flip(phi[:,:,-2:],axis=-1)], axis=-1)
    for a,b,p in zip(i,j,prob):
        if b!=a:
            name=tmpdir.name+"/%d.%d_phi.txt"%(a+1,b+1)
            with open(name, "w") as f:
                f.write('x_axis'+'\t%.3f'*nbins%tuple(bins)+'\n')
                f.write('y_axis'+'\t%.3f'*nbins%tuple(phi[a,b])+'\n')
                f.close()
            rst_line = 'Angle CA %d CB %d CB %d SPLINE TAG %s 1.0 %.3f %.5f'%(a+1,a+1,b+1,name,1.0,ASTEP)
            rst['phi'].append([a,b,p,rst_line])
            #if a==0 and b==9:
            #    with open(name,'r') as f:
            #        print(f.read())

    print("phi restraints:   %d"%(len(rst['phi'])))
    '''

    return rst

def set_random_dihedral(pose):
    nres = pose.total_residue()
    for i in range(1, nres):
        phi,psi=random_dihedral()
        pose.set_phi(i,phi)
        pose.set_psi(i,psi)
        pose.set_omega(i,180)

    return(pose)


#pick phi/psi randomly from:
#-140  153 180 0.135 B
# -72  145 180 0.155 B
#-122  117 180 0.073 B
# -82  -14 180 0.122 A
# -61  -41 180 0.497 A
#  57   39 180 0.018 L
def random_dihedral():
    phi=0
    psi=0
    r=random.random()
    if(r<=0.135):
        phi=-140
        psi=153
    elif(r>0.135 and r<=0.29):
        phi=-72
        psi=145
    elif(r>0.29 and r<=0.363):
        phi=-122
        psi=117
    elif(r>0.363 and r<=0.485):
        phi=-82
        psi=-14
    elif(r>0.485 and r<=0.982):
        phi=-61
        psi=-41
    else:
        phi=57
        psi=39
    return(phi, psi)


def read_fasta(file):
    fasta=""
    with open(file, "r") as f:
        for line in f:
            if(line[0] == ">"):
                continue
            else:
                line=line.rstrip()
                fasta = fasta + line;
    return fasta


def remove_clash(scorefxn, mover, pose):
    for _ in range(0, 5):
        if float(scorefxn(pose)) < 10:
            break
        mover.apply(pose)


def add_rst(pose, rst, n,u,sep1, sep2, params, nogly=False):

    pcut=params['PCUT']
    seq = params['seq']

    array=[]

    if nogly==True:
        array += [line for a,b,p,line in rst['dist'] if abs(a-b)>=sep1 and abs(a-b)<sep2 and seq[a]!='G' and seq[b]!='G' and p>=pcut]
        if params['USE_ORIENT'] == True:
            array += [line for a,b,p,line in rst['omega'] if abs(a-b)>=sep1 and abs(a-b)<sep2 and seq[a]!='G' and seq[b]!='G' and p>=pcut+0.5] #0.5
            array += [line for a,b,p,line in rst['theta'] if abs(a-b)>=sep1 and abs(a-b)<sep2 and seq[a]!='G' and seq[b]!='G' and p>=pcut+0.5] #0.5
            array += [line for a,b,p,line in rst['phi'] if abs(a-b)>=sep1 and abs(a-b)<sep2 and seq[a]!='G' and seq[b]!='G' and p>=pcut+0.6] #0.6
    else:
        array += [line for a,b,p,line in rst['dist'] if abs(a-b)>=sep1 and abs(a-b)<sep2 and p>=pcut]
        #print("p???",rst['dist'])
        #if params['USE_ORIENT'] == True:
            #array += [line for a,b,p,line in rst['omega'] if abs(a-b)>=sep1 and abs(a-b)<sep2 and p>=pcut+0.5]
            #array += [line for a,b,p,line in rst['theta'] if abs(a-b)>=sep1 and abs(a-b)<sep2 and p>=pcut+0.5]
            #array += [line for a,b,p,line in rst['phi'] if abs(a-b)>=sep1 and abs(a-b)<sep2 and p>=pcut+0.6] #0.6


    if len(array) < 1:
        return

    random.shuffle(array)

    # save to file
    tmpname = params['TDIR']+'/'+str(n)+'_'+str(u)+'minimize.cst'
    with open(tmpname,'w') as f:
        for line in array:
            f.write(line+'\n')
        f.close()

    # add to pose
    constraints = rosetta.protocols.constraint_movers.ConstraintSetMover()
    constraints.constraint_file(tmpname)
    constraints.add_constraints(True)
    constraints.apply(pose)

    os.remove(tmpname)
```
