## Supplementary material for "Predicting the alternative conformation of a known protein structure based on the distance map of AlphaFold2": codes and demos: README

###Procedure for (I) extracting alternative signals from AF2's prediction & (II) predicting the alternative conformation with Rosetta

I. Extracting alternative signals from DM based on the known structure

1) Store AlphaFold2 prediction results (https://colab.research.google.com/github/sokrypton/ColabFold/blob/main/AlphaFold2.ipynb) in the 'AF2_DM' folder. Sample data for Adenylate Kinase (apo:4AKE_B; holo:2ECK_B) is provided.
2) Place the known structure file (.cif format preferred) in the 'pdb' folder.
3) Execute the 'Step1-AlternativeSignalsAnalysis.py' script with the following arguments (detailed discription could be found with [-h]):
1: PDBid, e.g., 2ECK; 2: protein chain ID, e.g., B; 3: name of the .pickle file containing the predicted DM, e.g. AK_rec0_msaAll_27f14_all_rank_005_alphafold2_ptm_model_2_seed_000.pickle; 4 and 5: number of missing residues at the start and end in the initial structure(if applicable, default 0)

E.g., test the prepared data for Adenylate kinase with
>>python Step1-AlternativeSignalsAnalysis.py 2ECK B AK_rec0_msaAll_27f14_all_rank_005_alphafold2_ptm_model_2_seed_000.pickle 0 0 

Output files (saved in the 'distfile' folder):
1) Identified residue pairs: 'alt-name_xx.csv'
2) Alternative distance map: 'xx_altDM.npy'
3) Residue pair indices for alternative signals: 'xx_flag.npy'
4) Comentropy for predicted DM: 'Comentropy_xx.npy'
5) Signal number parameters: 'p-parameters.txt'

II. Predicting Alternative Conformations with Rosetta

Prerequisites: Install PyRosetta3 (http://www.pyrosetta.org/dow/pyrosetta3-download).
1) Prepare the initial known state structure (e.g. 2ECK.pdb) and its sequence file (AK.fasta) in the working directory. If there are ligands or ions in original structure file, please prepare with a clean protein file and make sure the sequence file matches the complete structure.
2) Run the 'Step2-Modeling.py' script with two arguments: 1-Path to the FASTA file; 2-Output structure name prefix

E.g., test the prediction for Adenylate kinase from holo to apo state
>>python Step2-Modeling.py ./fasta/AK.fasta ./results/alt-2ECK_n


Predicted models will be saved in the 'results' folder. The first 10 sampled models are offered for instance.
For more detailed information on the modeling process, refer to the "Materials and Methods: Model Building and Assessment" section of the associated paper.

Jiaxuan Li 
2024.09 
