## Supplementary material for "Predicting the alternative conformation of a known protein structure based on the distance map of AlphaFold2": codes and demos: Step1-AlternativeSignalsAnalysis.py

```
import pickle
import argparse
import json
import pandas as pd
from fitter import Fitter
from scipy.signal import argrelextrema,argrelmax,argrelmin
import numpy as np
np.set_printoptions(threshold=np.inf)
import matplotlib.pyplot as plt
from scipy.optimize import curve_fit
import gemmi
import torch
import math
from Bio.PDB import DSSP,PDBParser,MMCIFParser
import seaborn as sns

parser = argparse.ArgumentParser(description = 'analysis signals in AF2_DM')
parser.add_argument('PDBid',type = str, help = 'name of protein structure file for analysis')
parser.add_argument('ChainID',type = str, help = 'protein chain ID')
parser.add_argument('targetDM',type = str, help = 'name of AF2 predicted DM---.pickle file')
parser.add_argument('gap_beg',type = int, default=0, help = 'how many residues have gap in the begining of structure file with fasta (sequence used in the prediction of DM)')
parser.add_argument('gap_end',type = int, default=0, help = 'how many residues have gap in the end of structure file with fasta (sequence used in the prediction of DM)')
args = parser.parse_args()
pdbid = args.PDBid
chainid = args.ChainID
DMname = args.targetDM

def gauss1(x,a,u,sig):
    return a*np.exp(-(x-u)**2/(2*sig**2))/(sig*math.sqrt(2*math.pi))

def gaussian2(x,a1,a2,u1,u2,sig1,sig2):
    return a1*np.exp(-(x-u1)**2/(2*sig1**2))/(sig1*math.sqrt(2*math.pi))+a2*np.exp(-(x-u2)**2/(2*sig2**2))/(sig2*math.sqrt(2*math.pi))

def get_coords(chain, atoms = ('CB',), stack: bool = True, **kwargs):
    if stack:
        ret = np.array([[res[atom][0].pos.tolist() if not (atom == 'CB' and res.name == 'GLY') else res['CA'][0].pos.tolist() for atom in atoms] for res in chain], dtype=np.float32, **kwargs)
        return ret
    else:
        return np.array([res[atom][0].pos.tolist() if not (atom == 'CB' and res.name == 'GLY') else res['CA'][0].pos.tolist() for res in chain for atom in atoms], dtype=np.float32, **kwargs)

def get_cb_coords(chain, **kwargs):
    return np.array([get_real_or_virtual_CB(res) for res in chain], dtype=np.float32, **kwargs)

def get_real_or_virtual_CB(res, gly_ca: bool = True):
    if res.name != 'GLY':
        try:
            return res['CB'][0].pos.tolist()
        except Exception:
            pass
        #    LOGGER.debug(f"no CB for {res}")
    else:
        if gly_ca:
            #try:
            return res['CA'][0].pos.tolist()
            #except Exception as e:
            #    LOGGER.error(f"no CA for {res}")
            #    raise e
    #try:
    Ca = res['CA'][0].pos
    b = Ca - res['N'][0].pos
    c = res['C'][0].pos - Ca
    #except Exception as e:
    #    LOGGER.error(f"no enough anchor atoms for {res}")
    #    raise e
    a = b.cross(c)
    return (-0.58273431*a + 0.56802827*b - 0.54067466*c + Ca).tolist()

def prepare(pdb_id,chain_id,model_id,beg,end):
    st = gemmi.read_structure(f"{pdb_id}")
    p = MMCIFParser()
    #p = PDBParser() ##for PDB file
    structure = p.get_structure("Model",pdb_id)
    dssp = DSSP(structure[0],pdb_id)
    ss = []
    for row in dssp:
        ss.append(list(row[0:3]))
    chain = list(st[model_id][chain_id].get_polymer().first_conformer())[beg:end]
    three_letter_seq = [aa.name for aa in chain]
    one_letter_seq = gemmi.one_letter_code(three_letter_seq)
    #bb_coords = get_coords(chain, atoms=('CB',), stack=False)
    bb_coords = get_cb_coords(chain)
    return bb_coords, one_letter_seq, ss

def pkl_read(pkl_file, encoding='bytes'):
    print('Loading from: ' + pkl_file)
    fp = open(pkl_file, 'rb')
    contents = pickle.load(fp, encoding=encoding)
    fp.close()
    return contents
    
def find_peaks(d0,breadths,probs,D,peak_num):##d0_apo,breadth for ij,probs[i][j]; outputs:D_apo,peak_num_apo
    peak = []
    mark = 0
    breadth_range = breadths
    if d0>21.6875:
        if 'last' in breadth_range:
            D.append('True')
            peak_num.append('last')
        else:   
            D.append('False')
            peak_num.append("none")    
    else:
        if 'last' in breadth_range:
            breadth_range.remove('last')
            mark = 1
        for k in breadth_range:
            if d0 >= k[0] and d0 <= k[-1]:
                D.append('True')
                peak_num.append(breadth_range.index(k)+1)
                #print(k)
                break
            else:
                peak.append(1)
        if len(peak)==len(breadth_range):
            D.append('False')
            peak_num.append('none')
    if mark ==1:
        breadth_range.append('last')
    #print("findpeaks:",D[-1],peak_num[-1])
    return D,peak_num
def hist_pre(probs,x_bins):
    y_probs = []
    num_bins = 0
    for i in range(len(x_bins)):
        y_probs.extend(int(probs[i]/sum(probs)*100)*[x_bins[i]])
        num_bins += int(probs[i]/sum(probs)*100)
    print(y_probs,num_bins)
    return np.array(y_probs),num_bins

f = open('./AF2_DM/'+DMname,'rb')

name = 'alt-'+pdbid
data = pickle.load(f)
DM = data['distogram']['logits']
DM_logits = torch.from_numpy(DM).to(torch.float)
probs = torch.nn.functional.softmax(DM_logits,dim=-1)
bin_edges = data['distogram']['bin_edges']
bins = torch.from_numpy(bin_edges).to(torch.float)

##sequence begin?end?
seq_beg = args.gap_beg ##default = 0
seq_end = -args.gap_end ##default = None
if seq_beg >0 or seq_end <0:
    probs = probs[seq_beg:seq_end,seq_beg:seq_end,:]
probs = np.array(probs)
#probs = np.delete(probs,np.s_[res1:res2:1],0) ##if there are missing residues or gaps between sequence and structures
#probs = np.delete(probs,np.s_[res1:res2:1],1)
print(DM.shape,type(probs), "Length of protein:",probs.shape,len(bin_edges))

probs_apo = np.array(probs)
probs_holo = np.array(probs)
print(type(probs_apo))
Comentropy = np.zeros((probs.shape[0],probs.shape[0]))
L = probs.shape[0]
bins_64 = np.append(bin_edges,float(22.0))
x2 = bins_64
print(bins_64,len(bins_64),bins,'\n')
s = 0
Type = []
D_apo = []
D_holo = []
pairs = []
N_max = []
D0_apo = []
D0_holo = []
Breadth = []
U = []
n_contact = 0
n_last_apoflag = 0
n_last_holoflag = 0
n_mp = 0
n1p_apoT = 0
n1p_holoT =0
n1p_apoF = 0
n1p_holoF = 0
n_63 = 0
peak_num_apo = []
peak_num_holo = []
pdb_apo = pdbid
apo_flag = np.zeros((L,L))
holo_flag = np.zeros((L,L))
print("init apo/holo flags:", apo_flag.shape,holo_flag.shape)
#structure input
bb_coords_apo, one_letter_seq_apo,ss_apo = prepare('../pdb/'+pdb_apo+'.cif', chainid,0,0,None) 
print("APO",bb_coords_apo.shape,ss_apo[0],one_letter_seq_apo)
N_diffpeak = 0
N_samepeak = 0
N_aFhT = 0
N_hFaT = 0
N_FF=0

for i in range(0,L-6):
    for j in range(i+6,L):
        mean = sum((bins_64[x]-0.3125/2)*probs[i][j][x] for x in range(0,probs.shape[-1]))
        sigma = math.sqrt(sum(probs[i][j][x]*((bins_64[x]-0.3125/2-mean)**2) for x in range(0,probs.shape[-1])))       

        atom1_apo = bb_coords_apo[i]
        atom2_apo = bb_coords_apo[j]
        d_sub = [(atom1_apo[x]-atom2_apo[x])**2 for x in range(0,len(atom1_apo))]
        d0_apo = math.sqrt(sum(d_sub))
        #atom1_holo = bb_coords_holo[i]
        #atom2_holo = bb_coords_holo[j]
        #d_sub = [(atom1_holo[x]-atom2_holo[x])**2 for x in range(0,len(atom1_holo))]
        #d0_holo = math.sqrt(sum(d_sub))

        H = -sum(probs[i,j]*np.log(probs[i,j]))
        #print("Comentropy",i+1,j+1,H)
        Comentropy[i,j] = H
        p_contact = sum(probs[i][j][:19])
        if p_contact >0.5:
            n_contact+=1
            #print("contact p:",i+1,j+1,p_contact)
        if i in range(0,L) and j in range(0,L):
            n_max = 0
            pij = []
            breadth = []
            uu = []
            type = []
            maxvector = argrelextrema(np.array(probs[i][j][:-1]),np.greater)[0]
            minvector = argrelmin(np.array(probs[i][j][:-1]))[0]
            minv = []
            maxv = []
            #print("original max,min:",i+1,j+1,maxvector,minvector,len(maxvector),len(minvector))
            
            if (len(maxvector) <1) or (len(minvector) < 1):
                continue
            if float(max(probs[i,j,maxvector]))<1e-4:
                continue
            elif (len(maxvector)==1) and probs[i][j][-1]<=0.4*max(probs[i][j]):
                continue
            elif len(maxvector)>=2:
                for l in range(0,len(maxvector)-1):
                    if maxvector[l+1]-maxvector[l]>2 and probs[i][j][maxvector[l]]>1e-4:
                        maxv.append(maxvector[l])
                    else:
                        continue
                if maxvector[-1]-maxvector[-2]>2:
                    maxv.append(maxvector[-1])
                if len(maxv)>=2:
                    tag = 0
                    p_peaks = list(np.array(probs[i][j][maxv]))
                    p_max1 = max(p_peaks)
                    if sum(p_peaks)<=0.1:
                        continue ##all p_peak_max too low
                    index_max1 = p_peaks.index(p_max1)

                    for x in maxv:
                        if x>=57:
                            tag+=1
                    
                    u1 = bins_64[maxv[index_max1]]
                    #print("max1",maxvector,maxv,p_peaks,p_max1,index_max1,maxv[index_max1],u1)
                    p_peaks[index_max1]=0 ###max1=0,seek for second max
                    p_max2 = max(p_peaks)
                    index_max2 = p_peaks.index(p_max2)
                    u2 = bins_64[maxv[index_max2]]
                    if p_max2 <0.1*p_max1:
                        if float(probs[i][j][-1])>3*p_max1 and float(probs[i][j][-1])>0.1 and float(probs[i][j][-1])<0.8:
                            breadth.append("last")
                            uu.append(23.0)
                            n_max +=1
                            type.append("p2weak_last")
                            ##gaussian1 fit for 1 main peak
                            popt_solo,pcov_solo = curve_fit(gauss1,x2[:-1],np.array(probs[i][j][:-1]),maxfev=1000000,p0=[0.1,u1,1])
                            x_bin = np.arange(x2[0],x2[-1],0.1)
                            if popt_solo[1]>=20.75 or popt_solo[2]>3:
                                continue
                            #print("Gauss1:",popt_solo)
                            y_solo = [gauss1(xx,popt_solo[0],popt_solo[1],popt_solo[2]) for xx in x_bin]
                            fit_maxv_solo = argrelextrema(np.array(y_solo),np.greater)[0]
                            if max(y_solo)/float(probs[i][j][-1]) < 0.1:
                                continue
                            ll_solo = round(max(2.3125,popt_solo[1]-3*popt_solo[2]),4)
                            rr_solo = round(min(popt_solo[1]+3*popt_solo[2],21.6875),4)
                            if popt_solo[1]<=20.125:
                                n_max+=1
                                breadth.append((ll_solo,rr_solo))
                                uu.append(popt_solo[1])
                                type.append("p2weak_solofit")
                                #print("len(maxv)>=2, p2toolow only p1_maxv but last bin useful",i+1,j+1,p_max1,float(probs[i][j][-1]),ll_solo,rr_solo)
                        else:   
                            continue ###only 1 high peak berfore bins57
                        pass
                    elif (p_max1-p_max2)/p_max1<0.05 and (u2-u1)<2 and len(maxv)>=3:
                        #print("a.3 2maxv close",i+1,j+1,p_max1,p_max2)
                        p_peaks[index_max2]=0
                        p_max2 = max(p_peaks)
                        index_max2 = p_peaks.index(p_max2)
                        u2 = bins_64[maxv[index_max2]]
                    if p_max2 >0.1*p_max1:
                        #print(i+1,j+1,maxv,"probs",p_max1,p_max2,"uuuuuuuuuu",u1,u2)
                        popt,pcov = curve_fit(gaussian2,x2[:-1],np.array(probs[i][j][:-1]),maxfev=1000000,p0=[0.1,0.1,u1,u2,1,1])
                        x_bin = np.arange(x2[0],x2[-1],0.1)
                        u1_fit = min(popt[2],popt[3])
                        u2_fit = max(popt[2],popt[3])
                        #print("Params for Gaussian2:",popt,"fitted u",u1_fit,u2_fit)
                        if (popt[-1]>3 or popt[-1]<-3) and (popt[-2]>3 or popt[-2]<-3): ##两个峰展宽都大于3
                            #print("b.1 too wide",i+1,j+1)
                            continue
                        if u1_fit >20.125 and u2_fit >20.125:
                            #print("b.2 fitted too far",i+1,j+1)
                            continue
                        y = [gaussian2(xx,popt[0],popt[1],popt[2],popt[3],popt[4],popt[5]) for xx in x_bin]
                        fit_maxv = argrelextrema(np.array(y),np.greater)[0]
                        fit_minv = argrelmin(np.array(y))[0]
                        y_maxv = []
                        for e in fit_maxv:
                            y_maxv.append(y[int(e)])
                        if len(fit_maxv)>1 and len(fit_minv)>0:
                            #print("Fitted",i+1,j+1,"max_values",y_maxv,"min_values",y[int(fit_minv[0])])
                            y_minv = y[int(fit_minv[0])] 
                            mid = round(float(x_bin[fit_minv]),4)
                            if popt[2]<popt[3]:
                                ll_1 = round(max(2.3125,popt[2]-3*popt[2+2]),4)
                                rr_2 = round(min(popt[3]+3*popt[3+2],20.125),4)
                            else: 
                                ll_1 = round(max(2.3125,popt[3]-3*popt[3+2]),4)
                                rr_2 = round(min(popt[2]+3*popt[2+2],20.125),4)
                                #print(popt[2]+3*popt[2+2])
                            if abs(min(y_maxv)-max(y_minv,0))/min(y_maxv)<0.05: ##platform not 2 peaks!!!
                                
                                if max(y_maxv)/float(probs[i][j][-1]) > 0.1:
                                    n_max+=1
                                    breadth.append((ll_1,rr_2))
                                    uu.append(float((popt[2]+popt[3])/2))
                                    type.append("2p_platform_fit1")
                                #print("yyyyyyyyyyyyyyyyyyyy",y_maxv,y_minv)
                        
                            elif x_bin[fit_maxv[1]]-x_bin[fit_maxv[0]]>=1.5:
                                n_max +=2
                                breadth.append((ll_1,mid))
                                uu.append(min(popt[2],popt[3]))
                                type.append("2p_2fit_1")
                                breadth.append((mid,rr_2))
                                uu.append(max(popt[2],popt[3]))
                                type.append("2p_2fit_2")
                            elif max(y_maxv)/float(probs[i][j][-1]) > 0.1:
                                n_max+=1
                                breadth.append((ll_1,rr_2))
                                uu.append((popt[2]+popt[3])/2)
                                type.append("2p_2fit_restin1p")

                        else: 
                            if len(fit_maxv)==1 and max(y_maxv)/float(probs[i][j][-1])>0.1:
                                n_max += 1
                                vector = np.where(np.array(y)>0.005)[0]
                                breadth.append((round(x_bin[vector[0]],4),min(round(x_bin[vector[-1]],4),20.125)))
                                uu.append(float(x_bin[fit_maxv]))
                                type.append("2p_2fit_in1_over0.005")
                            #print("-----2 set of parameters used------------1 peak fitted---------------")

                    #### last bin counting!!!
                        if float(probs[i][j][-1])>0.1 and (sum(probs[i][j][57:63])<max(probs[i][j])/3):
                            print("d.1???",i+1,j+1)
                        if len(breadth)>=1 and max(breadth[:][-1])<20.125 and float(probs[i][j][-1])>0.1 and (sum(probs[i][j][57:63])>max(probs[i][j])/3):
                            breadth.append((20.125,21.6875))
                            uu.append(21.0)
                            n_max+=1
                            type.append("2p_bins57-63")
                        if float(probs[i][j][-1])>0.8*max(probs[i][j]) and float(probs[i][j][-1])>0.1 and float(probs[i][j][-1])<0.8:
                            n_max+=1
                            breadth.append('last')
                            uu.append(23.0)
                            type.append("2p_last")
                elif len(maxv)==1: #有效极大值点就一个
                    u1 = float(bins_64[maxv])
                    p_peak = sum(np.array(probs[i][j][int(maxv[0])-1:int(maxv[0])+2]))
                    if p_peak<0.1:
                        continue
                    ##gaussian1 fit for 1 main peak
                    popt_solo,pcov_solo = curve_fit(gauss1,x2[:-1],np.array(probs[i][j][:-1]),maxfev=1000000,p0=[0.1,u1,1])
                    x_bin = np.arange(x2[0],x2[-1],0.1)
                    if popt_solo[1]>=20.75 or popt_solo[2]>3: 
                        continue
                    y_solo = [gauss1(xx,popt_solo[0],popt_solo[1],popt_solo[2]) for xx in x_bin]
                    fit_maxv_solo = argrelextrema(np.array(y_solo),np.greater)[0]
                    if max(y_solo)/float(probs[i][j][-1])<0.1:
                        continue
                    ll_solo = round(max(2.3125,popt_solo[1]-3*popt_solo[2]),4)
                    rr_solo = round(min(popt_solo[1]+3*popt_solo[2],21.6875),4)
                    if float(probs[i][j][-1])>0.4 and float(probs[i][j][-1])<0.8 and p_peak>0.1:
                        n_max+=1
                        breadth.append((ll_solo,rr_solo))
                        uu.append(popt_solo[1])
                        type.append("1pmaxv=1_1fit")
                        print("ccccclen(maxv==1) 1_maxv but last bin useful",i+1,j+1,p_peak,float(probs[i][j][-1]),ll_solo,rr_solo)
                    elif (popt_solo[1]+popt_solo[2])<21.6875 and p_peak>0.1: ##last bin 进入u+1*sigma范围，不好区分最后一列是否是单独一个峰且是的化变化距离很小？
                        n_max+=1
                        breadth.append((ll_solo,rr_solo))
                        uu.append(popt_solo[1])
                        type.append("1pmaxv=1_1fit_u+1sigma")
                    if float(probs[i][j][-1])>p_peak and float(probs[i][j][-1])>0.1 and float(probs[i][j][-1])<0.8:
                        #print("LLLLLLLLLLLLen(maxv)==1, and last bin effective!!!!!!!",i+1,j+1)
                        breadth.append('last')
                        uu.append(23.0)
                        n_max +=1
                        type.append("1pmaxv=1_1fit_last")
                        #print("1_maxv but last bin useful",p_peak,float(probs[i][j][-1]),breadth)
                    else:   
                        continue ###only 1 high peak
            elif len(maxvector)==1: ##初始就一个极大值点
                p_maxvector = float(probs[i][j][maxvector])
                p_peak = sum(np.array(probs[i][j][int(maxvector[0])-1:int(maxvector[0])+2]))
                if p_maxvector/float(probs[i][j][-1])>1 or p_maxvector/float(probs[i][j][-1])<0.1:###one large peak / mainly on lastbin
                    continue
                elif p_peak>0.1 and float(probs[i][j][-1])<0.8: ##!!!!!
                    u = float(bins_64[maxvector])
                    popt_solo,pcov_solo = curve_fit(gauss1,x2[:-1],np.array(probs[i][j][:-1]),maxfev=1000000,p0=[0.1,u,1])
                    if popt_solo[1]>=20.75 or popt_solo[2]>3: ##拟合峰值所在距离不能与最后太近
                        continue
                    x_bin = np.arange(x2[0],x2[-1],0.1)
                    y_solo = [gauss1(xx,popt_solo[0],popt_solo[1],popt_solo[2]) for xx in x_bin]
                    fit_maxv_solo = argrelextrema(np.array(y_solo),np.greater)[0]
                    ll_solo = round(max(2.3125,popt_solo[1]-3*popt_solo[2]),4)
                    rr_solo = round(min(popt_solo[1]+3*popt_solo[2],21.6875),4)
                    if float(probs[i][j][-1])>0.4 and float(probs[i][j][-1])<0.8 and p_peak>0.1:
                        n_max+=1
                        breadth.append((ll_solo,rr_solo))
                        uu.append(popt_solo[1])
                        type.append("1pvector=1_1fit")
                        #print("LEN(MAXVECTOR==1) 1_maxv but last bin useful",p_maxvector,float(probs[i][j][-1]),ll_solo,rr_solo)
                    elif (popt_solo[1]+popt_solo[2])<21.6875 and p_peak>0.1: ##last bin 进入u+1*sigma范围，不好区分最后一列是否是单独一个峰且是的化变化距离很小？
                        n_max+=1
                        breadth.append((ll_solo,rr_solo))
                        uu.append(popt_solo[1])
                        type.append("1pvector=1_1fit_u+1sigma")
                        #print("11111sigma+u____LEN(MAXVECTOR==1) 1_maxv but last bin useful",p_maxvector,"lastbin",float(probs[i][j][-1]),ll_solo,rr_solo)
                    if float(probs[i][j][-1])>p_peak and float(probs[i][j][-1])>0.1 and float(probs[i][j][-1])<0.8:
                        n_max+=1
                        breadth.append('last')
                        uu.append(23.0)
                        type.append("1pvector=1_1fit_last")

            if n_max >=2:
                n_mp +=1
                D_apo,peak_num_apo = find_peaks(round(d0_apo,4),breadth,probs[i][j],D_apo,peak_num_apo)
                print(i+1,j+1,'breadth',breadth,"peaks",uu,"d0",round(d0_apo,4),'mark:',type,D_apo[-1],peak_num_apo[-1])
                pairs.append((i+1, j+1))
                N_max.append(n_max)
                Breadth.append(breadth)
                D0_apo.append(round(d0_apo,2))
                U.append(uu)
                if D_apo[-1] == 'True':
                    apo_flag[i,j]=1
                    if (peak_num_apo[-1] != 'last'):
                        if 'last' in breadth:
                            n_last_apoflag+=1
                        d_range = breadth[peak_num_apo[-1]-1]
                        ll = int((d_range[0]-2.3125)/0.3125)
                        rr = int((d_range[1]-2.3125)/0.3125)
                        probs_apo[i][j][ll:rr] = [0]*abs(ll-rr)
                        p_peak1 = sum(probs[i][j][ll:rr])
                        probs_apo[i][j] = (1/(1-p_peak1))*probs_apo[i][j]
                    elif peak_num_apo[-1] == 'last':
                        probs_apo[i][j][-1] = 0
                        probs_apo[i][j] = (1/(1-float(probs[i][j][-1])))*probs_apo[i][j]
            else: ##Single peak signal?
                probs_apo[i][j] = probs_apo[i][j]
                #probs_holo[i][j] = probs_holo[i][j]
                if n_max==1:
                    breadth = str(breadth[:]).replace("[","").replace("]","")
                    breadth = breadth.replace("'last'","22.0")
                    breadth = breadth.replace("(","").replace(")","")
                    breadth = breadth.split(',')
                    if breadth[0]=='22.0':
                        if d0_apo<21.6875:
                            n1p_apoT+=1
                            apo_flag[i,j]=1
                            n_last_apoflag+=1
                    else:
                        if (d0_apo<float(breadth[0]) or d0_apo>float(breadth[-1])):
                            n1p_apoT+=1 
                            apo_flag[i,j]=1
                            print("apoSINGLE",breadth,i+1,j+1,d0_apo)
maxh = np.max(Comentropy)
minh = np.min(Comentropy[np.where(Comentropy>0)])
#print("Counts:",n_mp,"Comentropy",maxh,minh)
for i in range(0,len(Comentropy)):
    for j in range(0,len(Comentropy[0])):
        Comentropy[i][j] = (maxh-Comentropy[i,j])/(maxh-minh)
#print("anti-normalized",np.min(Comentropy),np.max(Comentropy))
#print("count same/diffpeaks/ApoFalse/HoloFalse/FF:",N_samepeak,N_diffpeak,N_aFhT,N_hFaT,N_FF)
#print("Contact number and ratios:",n_contact,"apoflag",(len(pairs)-N_FF-N_aFhT)/n_contact,"holoflag",(len(pairs)-N_FF-N_hFaT)/n_contact)
print("1p numbers:",n1p_apoT,"multi-peak numbers",len(N_max))
print("proportion of 'last-bin' signals---p_last-bin:",round(n_last_apoflag/(len(N_max)+n1p_apoT),2))
print("density of 'alternative' signals---p_nl:",round((len(N_max)+n1p_apoT)/L,2))
with open('./distfile/p-parameters.txt','w')as f:
    f.write(name+'\n')
    f.write(str(round(n_last_apoflag/(len(N_max)+n1p_apoT),2))+'\n')
    f.write(str(round((len(N_max)+n1p_apoT)/L,2))) ##name, p_last-bin, p_nl

DM = pd.DataFrame({"res":pairs,
                   "n_max":N_max,
                   "d0_apo":D0_apo,
                   "apo_peak_loc":peak_num_apo,
                   "Apo_pred":D_apo,
                   "breadths":Breadth})
   
DM.to_csv('./distfile/'+name+'alternative signals_rec0_msaAll_rank005.csv')
np.save('./distfile/'+name+'_flag.npy',apo_flag)
#np.save('./distfile/'+name+'_holoflag.npy',holo_flag)
np.save('./distfile/'+name+'_altDM.npy',probs_apo)
#np.save('./distfile/'+name+'_altDM_holoflag.npy',probs_holo)
np.save('./distfile/Comentropy_'+name+'.npy',Comentropy)
```
