## Supplementary material for "Predicting the alternative conformation of a known protein structure based on the distance map of AlphaFold2": codes and demos: Step2-Modeling.py

```
import sys,os,json
import tempfile
import numpy as np

from arguments_AF2 import *
from dij_magH_noroll_Utils_cycle_ros_dist_npy import *
from pyrosetta import *
from pyrosetta.rosetta.protocols.minimization_packing import MinMover

os.environ["OPENBLAS_NUM_THREADS"] = "1"
# read and process parameters about alternative signals
with open('./distfile/p-parameters.txt','r')as f:
    lines=f.readlines()
    for line in lines:
        line=line.strip('\n')
    name=lines[0].strip('\n')
    pdbid=name.strip('alt-')
    p_last=float(lines[1])
    p_nl=float(lines[2])
    print("name",name,"p_last-bin",p_last,"p_nl",p_nl)

def main():
    ncycle = [1,2,3,4,5,6,7,8,9,10]
    U = [0.02,0.04,0.06,0.08,0.1,0.2,0.4,0.6,0.8,1.0,1.2,1.4,1.6,1.8,2.0]
    for n in ncycle:
        for u in U:
            print("ncycle U:",n,u)

            ########################################################
            # process inputs
            ########################################################

            # read params
            scriptdir = os.path.dirname(os.path.realpath(__file__))
            with open(scriptdir + '/data/params.json') as jsonfile:
                params = json.load(jsonfile)

            # get command line arguments
            args = get_args(params)
            print(args)

            # init PyRosetta
            init('-hb_cen_soft -relax:default_repeats 5 -default_max_cycles 200 -out:level 100')

            # Create temp folder to store all the restraints
            tmpdir = tempfile.TemporaryDirectory(prefix=args.wdir+'/')
            params['TDIR'] = tmpdir.name
            print('temp folder:     ', tmpdir.name)
            print("predict direction:",type)

            H = np.load('./distfile/Comentropy_'+name+'.npy')
            npz = np.load('./distfile/'+name+'_altDM.npy')
            dist_flag = np.load('./distfile/'+name+'_flag.npy')
            seq = read_fasta(args.FASTA)
            L = len(seq)
            params['seq'] = seq

            ########################################################
            # Scoring functions and movers
            ########################################################
            sf = ScoreFunction()
            sf.add_weights_from_file(scriptdir + '/data/scorefxn.wts')
            
            sf1 = ScoreFunction()
            sf1.add_weights_from_file(scriptdir + '/data/scorefxn1.wts')

            sf_vdw = ScoreFunction()
            sf_vdw.add_weights_from_file(scriptdir + '/data/scorefxn_vdw.wts')

            sf_cart = ScoreFunction()
            sf_cart.add_weights_from_file(scriptdir + '/data/scorefxn_cart.wts')

            mmap = MoveMap()
            mmap.set_bb(True)
            mmap.set_chi(True)
            mmap.set_jump(True)

            min_mover = MinMover(mmap, sf, 'lbfgs_armijo_nonmonotone', 0.0001, True)
            min_mover.max_iter(1000)

            min_mover1 = MinMover(mmap, sf1, 'lbfgs_armijo_nonmonotone', 0.0001, True)
            min_mover1.max_iter(1000)

            min_mover_vdw = MinMover(mmap, sf_vdw, 'lbfgs_armijo_nonmonotone', 0.0001, True)
            min_mover_vdw.max_iter(500)

            min_mover_cart = MinMover(mmap, sf_cart, 'lbfgs_armijo_nonmonotone', 0.0001, True)
            min_mover_cart.max_iter(1000)
            min_mover_cart.cartesian(True)

            repeat_mover = RepeatMover(min_mover, n)


            ########################################################
            # initialize pose
            ########################################################
            apo = pose_from_pdb("./pdb/"+pdbid+".pdb")  #AK_holo
            pose = Pose()
            pose.assign(apo)
            switch = SwitchResidueTypeSetMover("centroid")
            switch.apply(pose)

            # mutate GLY to ALA
            for i,a in enumerate(seq):
                if a == 'G':
                    mutator = rosetta.protocols.simple_moves.MutateResidue(i+1,'ALA')
                    mutator.apply(pose)
                    print('mutation: G%dA'%(i+1))

            #set_random_dihedral(pose)
            #remove_clash(sf_vdw, min_mover_vdw, pose)
            rst = gen_rst(npz,u,dist_flag,H,tmpdir,params,pose,p_last,p_nl)
            print("dist rst file write!")

            ########################################################
            # minimization
            ########################################################

            if args.mode == 0:

                # short
                print('short')
                add_rst(pose, rst, 1, 12, params)
                repeat_mover.apply(pose)
                min_mover_cart.apply(pose)
                remove_clash(sf_vdw, min_mover1, pose)

                # medium
                print('medium')
                add_rst(pose, rst, 12, 24, params)
                repeat_mover.apply(pose)
                min_mover_cart.apply(pose)
                remove_clash(sf_vdw, min_mover1, pose)

                # long
                print('long')
                add_rst(pose, rst, 24, len(seq), params)
                repeat_mover.apply(pose)
                min_mover_cart.apply(pose)
                remove_clash(sf_vdw, min_mover1, pose)

            elif args.mode == 1:

                # short + medium
                print('short + medium')
                add_rst(pose, rst, 3, 24, params)
                repeat_mover.apply(pose)
                min_mover_cart.apply(pose)
                remove_clash(sf_vdw, min_mover1, pose)

                # long
                print('long')
                add_rst(pose, rst, 24, len(seq), params)
                repeat_mover.apply(pose)
                min_mover_cart.apply(pose)
                remove_clash(sf_vdw, min_mover1, pose)

            elif args.mode == 2:

                # short + medium + long
                print('short + medium + long')
                add_rst(pose, rst,n,u, 1, len(seq), params)
                repeat_mover.apply(pose)
                min_mover_cart.apply(pose)
                remove_clash(sf_vdw, min_mover1, pose)

            ########################################################
            # full-atom refinement
            ########################################################

            if args.fastrelax == True:

                sf_fa = create_score_function('ref2015')
                sf_fa.set_weight(rosetta.core.scoring.atom_pair_constraint, 5)
                sf_fa.set_weight(rosetta.core.scoring.dihedral_constraint, 1)
                sf_fa.set_weight(rosetta.core.scoring.angle_constraint, 1)

                mmap = MoveMap()
                mmap.set_bb(True)
                mmap.set_chi(True)
                mmap.set_jump(True)

                relax = rosetta.protocols.relax.FastRelax()
                relax.set_scorefxn(sf_fa)
                relax.max_iter(1000)
                relax.dualspace(True)
                relax.set_movemap(mmap)

                pose.remove_constraints()
                switch = SwitchResidueTypeSetMover("fa_standard")
                switch.apply(pose)

                print('relax...')
                params['PCUT'] = 0.15
                add_rst(pose, rst,n,u, 1, len(seq), params, True)
                relax.apply(pose)

                # mutate ALA back to GLY
                for i,a in enumerate(seq):
                    if a == 'G':
                        mutator = rosetta.protocols.simple_moves.MutateResidue(i+1,'GLY')
                        mutator.apply(pose)
                        print('mutation: A%dG'%(i+1))

            ########################################################
            # save final model
            ########################################################
            pose.dump_pdb(args.OUT+str(n)+"_U"+str(u)+".pdb")


if __name__ == '__main__':
    main()
```
