## Supplementary figures and images for "Predicting the alternative conformation of a known protein structure based on the distance map of AlphaFold2"

### AK_rec0_msaAll_27f14_coverage.png

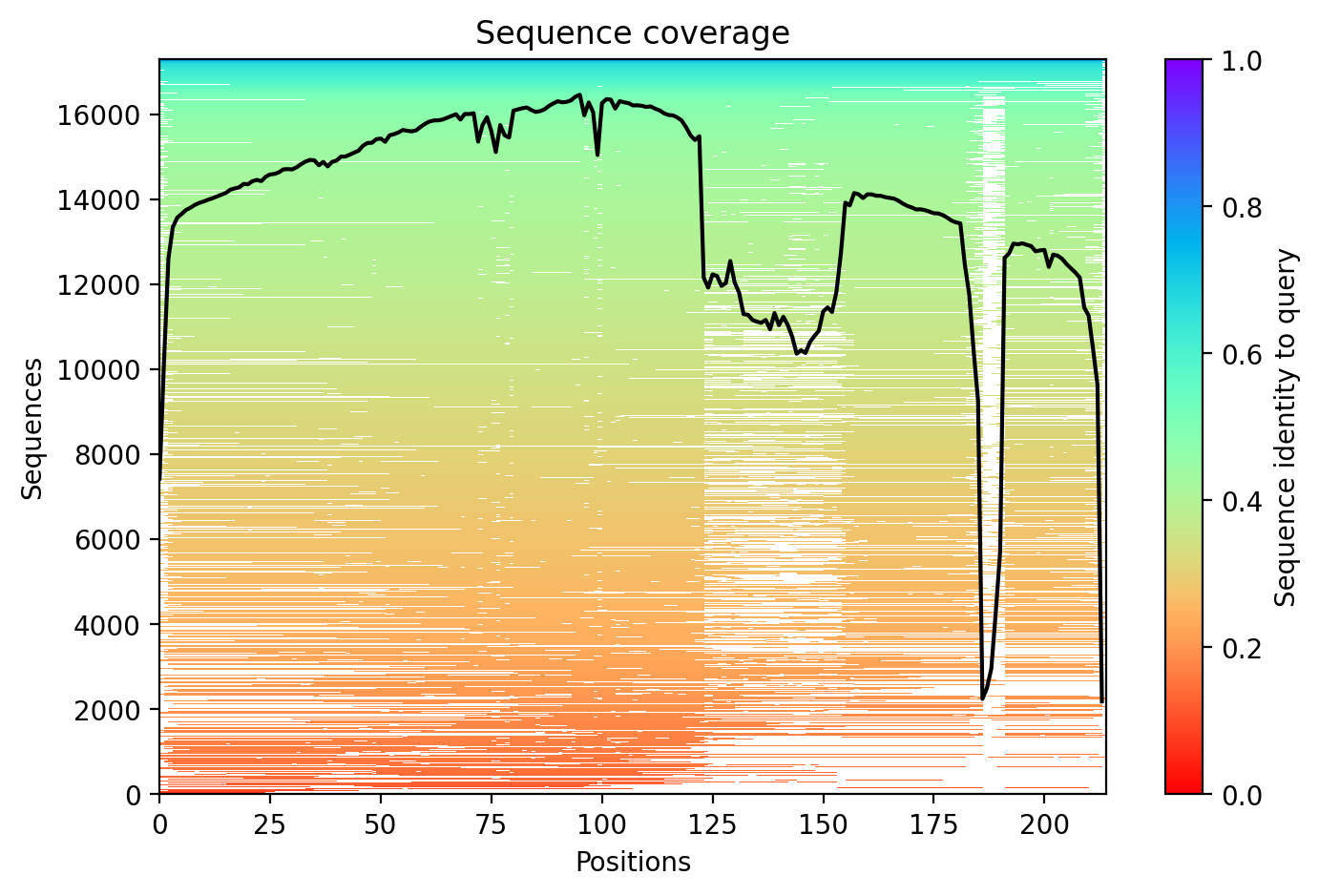

### AK_rec0_msaAll_27f14_pae.png

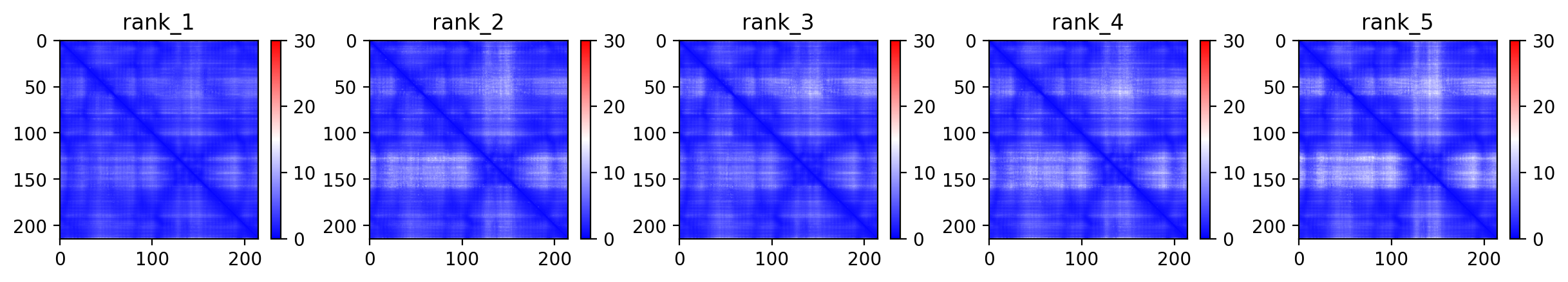

### AK_rec0_msaAll_27f14_plddt.png

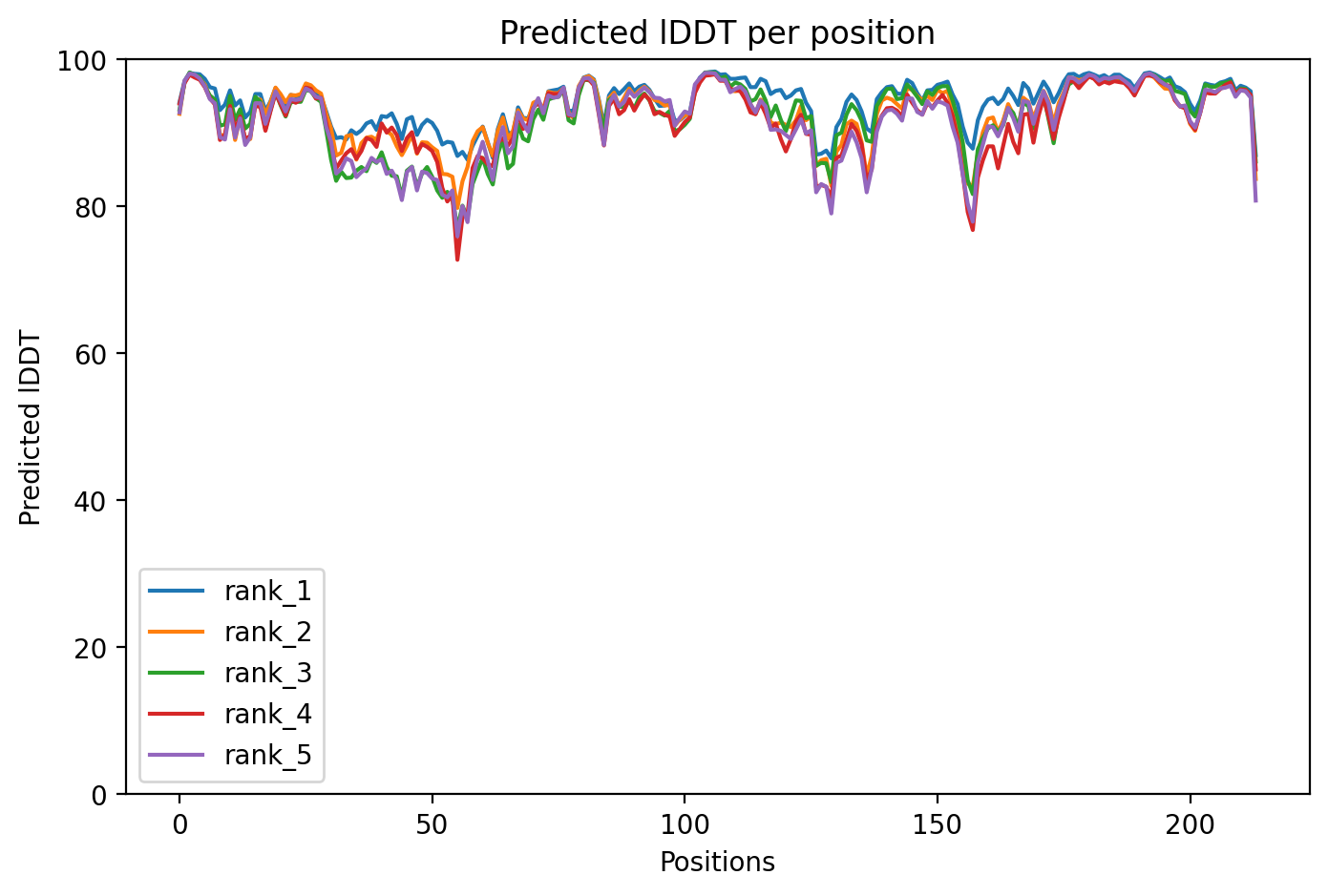
